# DONSON is required for CMG helicase assembly in the mammalian cell cycle

**DOI:** 10.1101/2023.08.16.553614

**Authors:** Cecile Evrin, Vanesa Alvarez, Johanna Ainsworth, Ryo Fujisawa, Constance Alabert, Karim P.M. Labib

## Abstract

DONSON is one of 13 genes mutated in a form of primordial microcephalic dwarfism known as Meier-Gorlin Syndrome. The other 12 encode components of the CDC45-MCM-GINS helicase, around which the eukaryotic replisome forms, or are factors required for helicase assembly during DNA replication initiation. A role for DONSON in CDC45-MCM-GINS assembly was unanticipated, since DNA replication initiation can be reconstituted *in vitro* with purified proteins from budding yeast, which lacks DONSON. Using mouse embryonic stem cells as a model for the mammalian helicase, we show that DONSON binds directly but transiently to CDC45-MCM-GINS during S-phase and is essential for chromosome duplication. Rapid depletion of DONSON leads to the disappearance of the CDC45-MCM-GINS helicase from S-phase cells and our data indicate that DONSON is dispensable for loading of the MCM2-7 helicase core onto chromatin during G1-phase, but instead is essential for CDC45-MCM-GINS assembly during S-phase. These data identify DONSON as a missing link in our understanding of mammalian chromosome duplication and provide a molecular explanation for why mutations in human DONSON are associated with Meier-Gorlin syndrome.

## Introduction

In eukaryotic cells, genome duplication is initiated at many origins of DNA replication along each chromosome by assembly of the CMG helicase (CDC45-MCM-GINS), which unwinds the parental DNA duplex at DNA replication forks (Bell & Labib, 2016; Costa & Diffley, 2022; Riera *et al*, 2017). CMG helicase assembly is highly regulated to ensure that cells make a single copy of every chromosome during each cell cycle. The helicase is assembled in two steps that are temporally separated from each other. Firstly, the six MCM2-7 ATPases are loaded at origins of DNA replication during G1-phase as rings around double-stranded DNA (dsDNA), by the Origin Recognition Complex (ORC) together with the CDC6 and CDT1 proteins, to form MCM2-7 double hexamers that lack helicase activity (Bell & Labib, 2016; Costa & Diffley, 2022; Riera *et al*., 2017). Subsequently, the CDC45 protein and the four-protein GINS complex are recruited by multiple CMG assembly factors to MCM2-7 double hexamers, in a process controlled by the CDC7 kinase and Cyclin Dependent Kinases. In this way, MCM2-7 double hexamers are converted into two nascent CMG helicases, around which two replisomes assemble at a pair of bi-directional replication forks. CMG remains stably associated with replication forks throughout elongation, until termination when the helicase is ubiquitylated and then disassembled by the p97 ATPase (Dewar & Walter, 2017; Maric *et al*, 2014; Moreno & Gambus, 2020; Sonneville *et al*, 2017; Villa *et al*, 2021).

The CMG helicase is essential for cell viability, but partial loss of function or defects in helicase assembly cause a form of microcephalic primordial dwarfism in humans called Meier Gorlin Syndrome, which is associated with hypomorphic mutations in 13 genes (Bellelli & Boulton, 2021; Klingseisen & Jackson, 2011; Nielsen-Dandoroff *et al*, 2023). Of these, twelve encode helicase subunits or factors with a well characterised role in CMG helicase assembly (three subunits of ORC, the CDC6 protein, CDT1 and its regulator Geminin, CDC45, three subunits of MCM2-7 and two components of GINS). The 13^th^ gene is called *DONSON* (Downstream Neighbor of SON), mutations in which lead not only to Meier Gorlin Syndrome (Reynolds *et al*, 2017) but also to other forms of microcephalic dwarfism with additional defects, such as Microcephaly with Short Stature and Limb Abnormalities (Schulz *et al*, 2018) and Microcephaly-Micromelia Syndrome (Evrony *et al*, 2017). Previous studies showed that depletion of human DONSON by RNA interference causes defects in genome stability (Fuchs *et al*, 2010) and mutation of the *Drosophila* orthologue causes DNA replication defects (Bandura *et al*, 2005). However, a role for DONSON in CMG helicase assembly was not anticipated, in contrast to other Meier Gorlin genes such as ORC, CDC6 and CDT1, since DONSON orthologues are absent from most fungal species including budding yeast, for which the process of CMG helicase assembly and DNA replication initiation can be reconstituted *in vitro* with a set of purified proteins that are conserved not only in fungi but also in animals and plants (Yeeles *et al*, 2015; Yeeles *et al*, 2017).

Human DONSON was found to co-purify with components of the CMG helicase and with the TIMELESS-TIPIN complex that binds to CMG in the replisome (Reynolds *et al*., 2017), suggesting a role for DONSON at DNA replication forks. Consistent with this view, replication forks stall more frequently after depletion of DONSON in HeLa cells (Reynolds *et al*., 2017). Moreover, DONSON is important for forks to traverse inter-strand DNA crosslinks (Zhang *et al*, 2020), indicating a role in replication-coupled DNA repair. However, it is unclear whether DONSON is recruited to forks under specific conditions or travels continuously with replication forks as part of the replisome. In previous work, DONSON was found to be enriched on newly replicated chromatin in some experiments (Alvarez *et al*, 2023; Reynolds *et al*., 2017) but not in others (Alabert *et al*, 2014; Alvarez *et al*., 2023; Nakamura *et al*, 2021; Wessel *et al*, 2019). Until now, therefore, the molecular function of DONSON during chromosome replication in mammalian cells has remained unclear.

We recently showed that mouse embryonic stem cells (mouse ES cells) provide an excellent model system for studying the regulation of the mammalian CMG helicase (Fujisawa *et al*, 2022; Villa *et al*., 2021), taking advantage of the fact that mouse ES cells proliferate quickly, spend much of their cell cycle in S-phase and are a rich source of CMG and replisomes. Using mouse ES cells and recombinant proteins as tools, we show here that mammalian DONSON interacts directly with components of the CMG helicase and is essential for CMG assembly during the S-phase of the cell cycle.

## Results & Discussion

### DONSON binds transiently to CMG helicase components during chromosome replication in mouse ES cells

To monitor the association of DONSON and the CMG helicase in mouse ES cells, we used CRISPR-Cas9 to introduce the TAP tag (Tandem Affinity Purification) into the *Psf1* and *Donson* loci, generating homozygous cell lines expressing functional tagged proteins (Fig EV1). Control mouse ES cells, or *TAP-Psf1* cells, were grown without further treatment, or in the presence of the p97 inhibitor CB-5083 that blocks CMG disassembly after DNA replication termination and causes the helicase to accumulate on chromatin with ubiquitylation of the MCM7 subunit of CMG (Villa *et al*., 2021). Cell extracts were treated with DNase to release proteins from chromatin and then incubated with immunoglobulin-coated magnetic beads (Fig 1A). This led to specific isolation of GINS and other CMG subunits from *TAP-Psf1* cells, together with partners of CMG such as DNA polymerase ε (Fig 1A, lanes 7-8). We found that DONSON also co-purified specifically with TAP-PSF1, consistent with previous observations for human DONSON (Reynolds *et al*., 2017; Zhang *et al*., 2020). Similarly, CMG components and CMG partner proteins co-purified specifically with TAP-tagged DONSON (Fig 1B, lanes 7-8). Together with previous data for human DONSON (Reynolds *et al*., 2017; Zhang *et al*., 2020), these findings indicated that mammalian DONSON associates with components of CMG, or with partners of the helicase.

**Figure 1.**
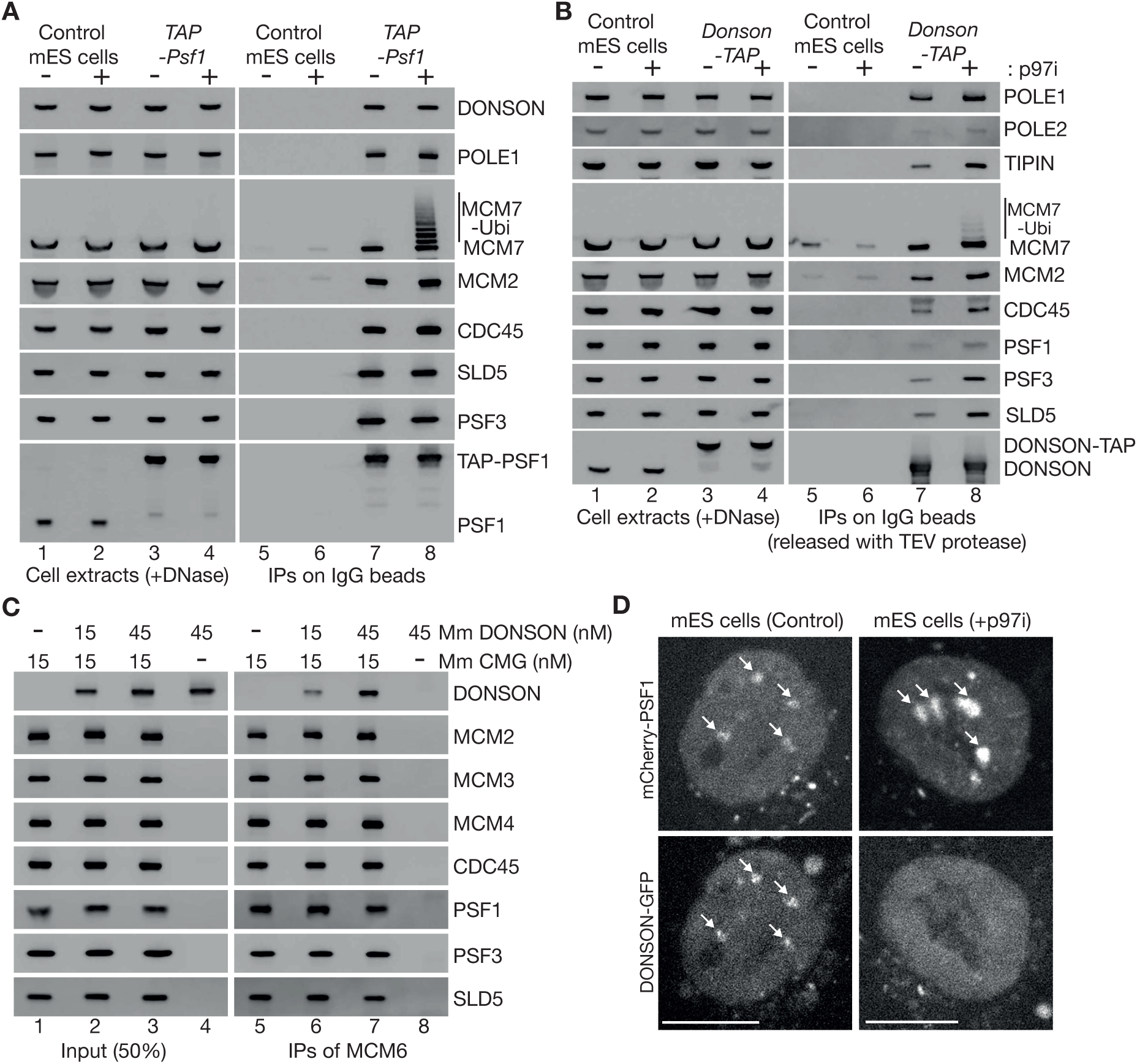
DONSON binds transiently to CMG helicase components during chromosome replication in mouse ES cells. **(A)** Extracts of control and *TAP-Psf1* cells were incubated with magnetic beads coated with IgG. The indicated proteins were detected by immunoblotting. **(B)** Similar experiment to (A) with control and *DONSON-TAP* cells. Proteins bound to beads were released by TEV cleavage, to avoid interference of the TAP tag in immunoblots for proteins of similar size to DONSON-TAP. **(C)** Recombinant mouse DONSON and CMG (Fig EV2) were purified and mixed as indicated (see Materials and Methods), before isolation of CMG on beads coated with antibodies to MCM6. **(D)** *mCherry-Psf1 Donson-GFP* mouse ES cells were untreated (left panels) or incubated for 3 hours with 5 µM CB-5083 (right panels), before imaging by spinning-disk confocal microscopy. Arrows indicate accumulation of mCherry-PSF1 and DONSON-GFP on heterochromatin patches. Scalebars correspond to 10 µm.

To test whether DONSON binds CMG directly, we expressed recombinant mouse DONSON in bacteria and generated haploid budding yeast cells that co-express the 11 subunits of mouse CMG (see Materials and Methods). After purification of recombinant mouse DONSON and CMG (Fig EV2), the two proteins were mixed and incubated *in vitro* (Figure 1C, lanes 1-4), before isolation of CMG on beads coated with antibodies to the MCM6 subunit of the helicase (Figure 1C, lanes 5-7). Association of DONSON with the beads was dependent upon the presence of CMG (Fig 1C, compare lanes 6-8), indicating that DONSON binds directly to components of the helicase.

We previously showed that the association of the CMG helicase with replicating chromatin can be monitored in mouse ES cells by spinning disk confocal microscopy, via condensed patches of constitutive heterochromatin (Fujisawa *et al*., 2022; Villa *et al*., 2021). CMG associates transiently with such patches when they replicate during S-phase, with similar timing to markers of DNA synthesis such as the polymerase processivity factor PCNA. However, if helicase disassembly is blocked by inhibition of p97, CMG remains on heterochromatin patches after replication, together with other proteins that are bound to CMG in the replisome, such as CTF4/WDHD1/AND-1, CLASPIN and DNA polymerase ε (Villa *et al*., 2021).

To monitor the association of DONSON with CMG on replicating chromatin, we generated mouse ES cells that expressed functional forms of mCherry-PSF-1 and DONSON-GFP from the endogenous loci (Fig EV1C-D; see Materials and Methods). In untreated cells, mCherry-PSF1 was present in the nucleus and accumulated on heterochromatin patches in 21% cells (Fig 1D, Control, n = 106 of 508 cells), consistent with past observations for the SLD5 subunit of GINS (Villa *et al*., 2021). Moreover, DONSON-GFP co-localised with mCherry-PSF1 in 91% of such cells (Fig 1D, Control, n = 96 of 106 cells with mCherry-PSF1 on heterochromatin patches), indicating that DONSON is present on replicating chromatin. We note that a previous study of human DONSON reported that co-localisation of DONSON with PSF1 was predominantly detected on euchromatin during early S-phase, using a proximity ligation assay (Zhang *et al*., 2020). We suggest that the greater density of active origins in euchromatin might facilitate detection in such assays, but our data show directly that DONSON also localises with PSF1 on replicating heterochromatin.

When cells were treated with p97 inhibitor to block CMG disassembly during DNA replication termination, mCherry-PSF-1 remained on heterochromatin patches in about a quarter of the cells (n = 130 of 491 cells), consistent with previous observations for GFP-SLD5 (Villa *et al*., 2021). However, DONSON-GFP was not detected on chromatin under such conditions (Fig 1D, p97i, DONSON-GFP failed to co-localise with mCherry-PSF1 in 98% of cells with mCherry-PSF1 on heterochromatin, n = 130). Therefore, although DONSON binds directly to components of the CMG helicase (Fig 1C) and is present on chromatin during replication in unperturbed cells (Fig 1D), its association with CMG is transient by comparison with replisome components such as CTF4, CLASPIN or DNA polymerase ε (Villa *et al*., 2021).

### DONSON is essential for DNA replication in mouse ES cells

As a first step towards exploring DONSON function in mouse ES cells, we used the Cas9-D10A ‘nickase’ and pairs of guide RNAs to introduce small deletions or insertions into both copies of exon 1 (Fig EV3A) or exon 4 (Fig EV4A) of the *Donson* gene. However, all tested clones still expressed DONSON protein, despite containing small deletions or insertions in both alleles (Fig EV3-4), which should have produced frameshifts in many cases. This indicates that natural selection favours the survival of *Donson* deletions that preserve protein expression and suggests that *Donson* is an essential gene in mouse ES cells, as is the case during early mouse development (Evrony *et al*., 2017). Therefore, we generated a *Donson* allele that allowed conditional degradation of DONSON protein, using CRISPR-Cas9 to integrate the BromoTag degron cassette (Bond *et al*, 2021) at the start of the DONSON coding sequence (Fig EV5). The BromoTag degron is recognised by heterobifunctional molecules known as ‘Proteolysis Targeting Chimeras’ (PROTACs), which simultaneously bind to the ubiquitin ligase CUL2^VHL^ (Bond *et al*., 2021). Addition of the AGB1 PROTAC did not affect the growth of control cells but blocked proliferation of *BromoTag-Donson* cells (Fig 2A), reflecting degradation of BromoTag-DONSON protein (Fig 2B) in response to ubiquitylation by CUL2^VHL^. This confirmed that DONSON is essential for viability in mouse ES cells.

**Figure 2.**
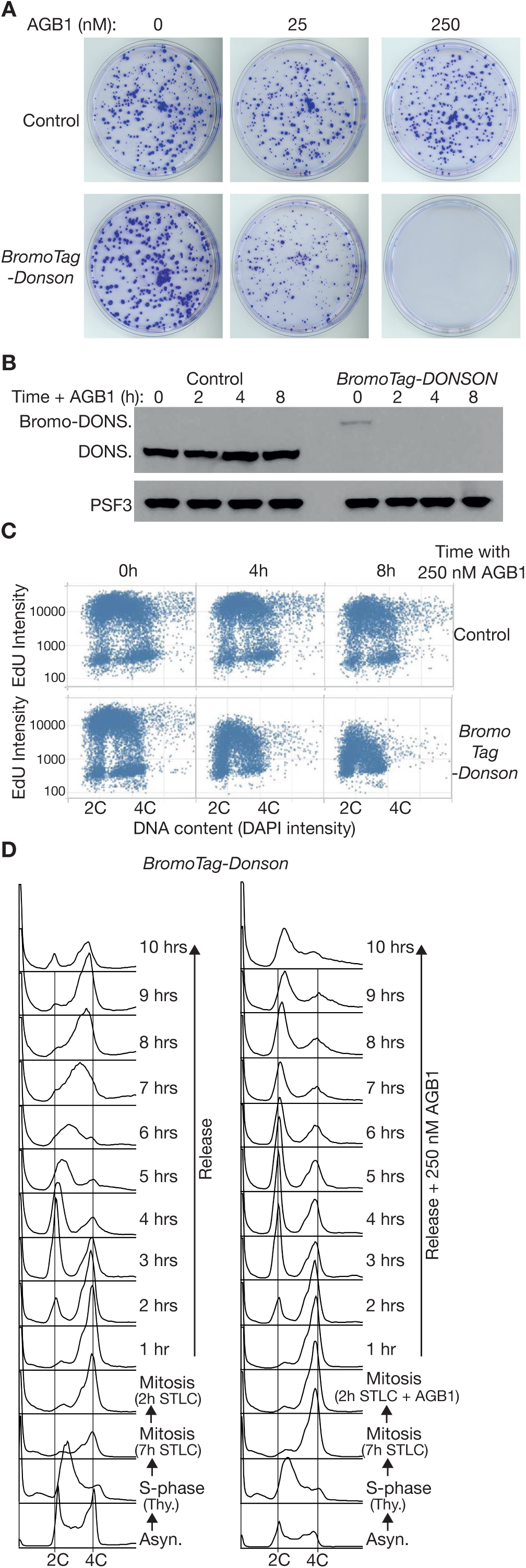
DONSON is essential for DNA replication in mouse embryonic stem cells. **(A)** Control and *BromoTag-Donson* cells were grown for 7 days in the presence of the indicated concentrations of AGB1, before staining with crystal violet solution as described in Materials and Methods. **(B)** Immunoblots showing the degradation of BromoTag-DONSON after treatment with 250 nM AGB1 for 0, 2, 4 or 8 hours. **(C)** The indicated cells were treated with 250 nM AGB1 as shown, before monitoring EdU incorporation into genomic DNA by Quantitative Image Based Cytometry (see Materials and Methods). **(D)** *BromoTag-Donson* cells were synchronised in early S-phase by incubation with 1.25 mM thymidine for 18 hours. Cells were then released into fresh medium containing 5 µM STLC for 9 hours, +/-250 nM AGB1 for the last two hours as indicated. Cells were then released into fresh medium lacking STLC. DNA content was monitored throughout the experiment by flow cytometry.

We then exposed asynchronous populations of control and *BromoTag-Donson* cells to AGB1 for different times, and at each point subjected cells to a short pulse of EdU. Incorporation of EdU into genomic DNA was monitored by Quantitative Image-Based Cytometry, which individually analyses thousands of cells and quantifies the intensity of their emitted fluorescence. EdU incorporation was reduced upon degradation of BromoTag-DONSON (Fig 2C), indicating that DONSON is required for DNA replication in the mammalian cell cycle.

To confirm that DONSON was specifically important for cell cycle progression during S-phase, we synchronised *BromoTag-Donson* cells in mitosis by addition of the kinesin inhibitor S-trityl-L-cysteine (STLC) after an initial thymidine arrest (Fig 2D and Materials and Methods), and then treated half of the culture with AGB1 for two hours. Degradation of BromoTag-DONSON did not prevent progression through mitosis and cell division, upon release into fresh medium lacking STLC, but instead blocked DNA replication during S-phase of the subsequent cell cycle (Fig 2D).

### DONSON is essential for CMG helicase assembly on loaded MCM2-7 complexes during S-phase of the mammalian cell cycle

The importance of DONSON for DNA replication could potentially indicate an essential role for DONSON in DNA synthesis at replication forks, or in progression of the CMG helicase. However, depletion of DONSON in human cells did not reduce the speed of replication fork progression (Reynolds *et al*., 2017) and recent work showed that a human replisome can be reconstituted *in vitro* with a set of purified proteins lacking DONSON, if the initiation step is bypassed by inclusion of recombinant CMG helicase (Baris *et al*, 2022). Moreover, the reconstituted human replisome supports DNA synthesis and fork progression at rates that approximate the fastest seen *in vivo*. Therefore, the ability of DONSON to bind components of the CMG helicase might instead indicate a role during assembly or activation of the CMG helicase in mammalian cells, even though helicase assembly and activation in budding yeast does not require DONSON and instead is mediated by other factors that have orthologues in metazoa (Yeeles *et al*., 2017).

As a first step towards testing whether DONSON might be required for CMG helicase assembly, *TAP-Psf1* and *BromoTag-Donson TAP-Psf1* cells were treated with AGB1, before isolation of CMG from cell extracts by immunoprecipitation of TAP-PSF1 (Fig 3A). Treatment of *TAP-Psf1* cells with AGB1 had no impact on the level of CMG (Fig 3A, lanes 7-9), whereas degradation of BromoTag-DONSON led to a reduction in the helicase without affecting the level of GINS (Fig 3A, lanes 10-12, compare the PSF3 & SLD5 subunits of GINS in TAP-PSF1 immunoprecipitates, with CDC45 and MCM2).

**Figure 3.**
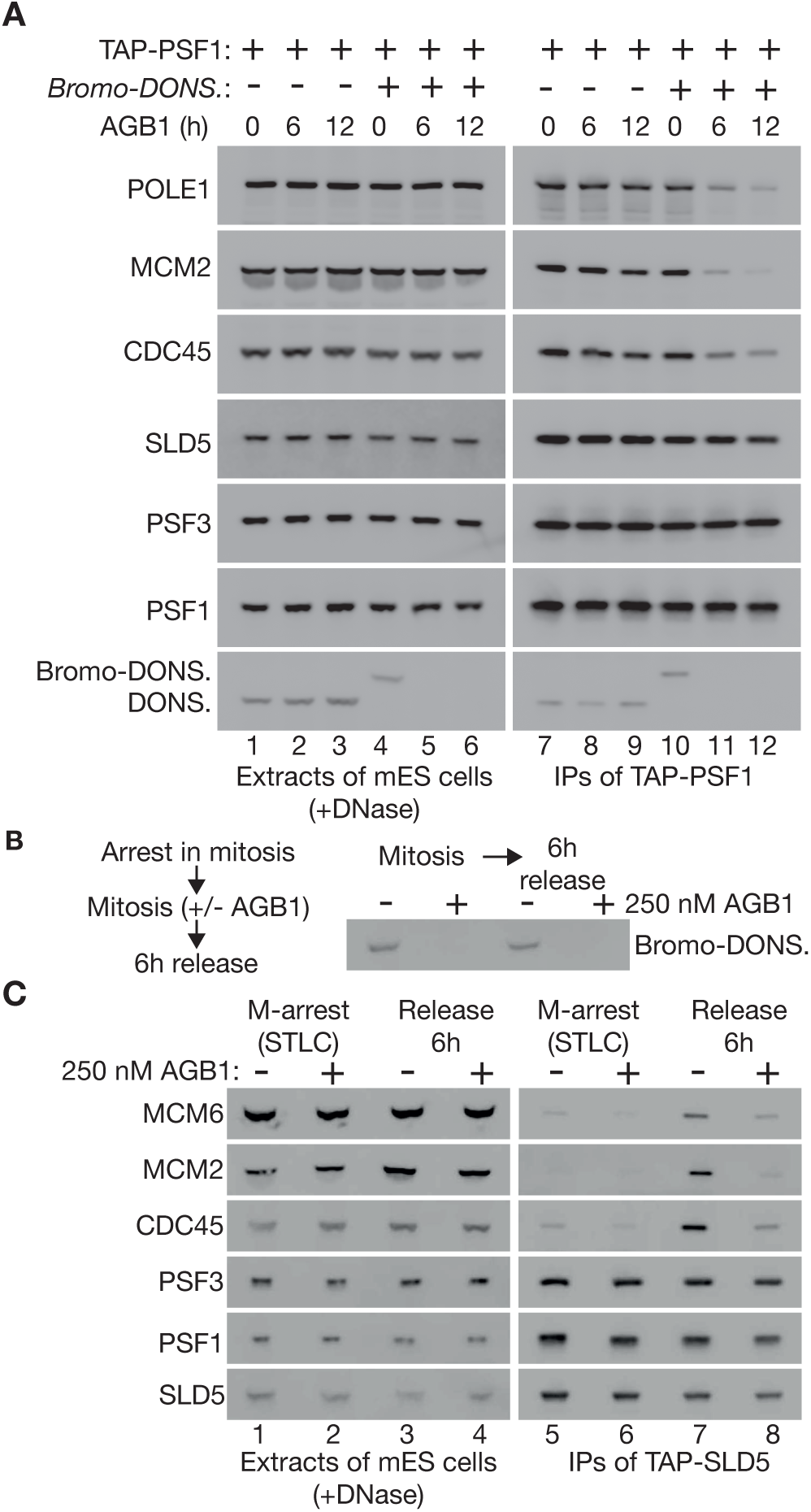
DONSON is required for presence of the CMG helicase in S-phase cells. **(A)** *TAP-Psf1* and *TAP-Psf1 BromoTag-Donson* cells were incubated with 250 nM AGB1 for the indicated times (h = hours), before isolation of TAP-PSF1 from cell extracts by immunoprecipitation on beads coated with IgG. The indicated factors were monitored by immunoblotting. **(B)** *TAP-SLD5 BromoTag-Donson* cells were treated as in Fig 2D and the level of BromoTag-DONSON was monitored at the indicated points. **(C)** Presence of the CMG helicase was monitored in the experiment in (B), in mitotic cells and upon release for 6 hours into S-phase, by immunoprecipitation of TAP-SLD5.

To monitor the appearance of the CMG helicase in S-phase cells, we synchronised *BromoTag-Donson TAP-Sld5* cells in mitosis as above (Fig 2D), before incubation for a further two hours plus or minus AGB1, followed by release for 6 hours into the subsequent cell cycle (Fig 3B), corresponding to mid-S-phase for cells containing DONSON (Fig 2D). Immunoprecipitation of the SLD5 subunit of GINS indicated that GINS was present throughout the experiment (Fig 3C, lanes 5-8, association of SLD5 with PSF1 and PSF3 subunits of GINS), whereas CMG was absent from mitotic cells as expected (Fig 3C, lanes 5-6, GINS did not associate with CDC45-MCM2-MCM6 subunits of CMG), was present in S-phase cells containing BromoTag-DONSON (Fig 3C, lane 7) but was much reduced in S-phase cells lacking BromoTag-DONSON (Fig 3C, lane 8). These findings indicate that DONSON is an important determinant of the level of the CMG helicase in the mammalian cell cycle.

The reduced level of CMG in the absence of DONSON could potentially be explained by DONSON preserving the integrity of the helicase at replication forks, but this seems unlikely for several reasons. As noted above, depletion of DONSON in human cells did not reduce the rate of fork progression (Reynolds *et al*., 2017). Moreover, recombinant mouse (Fig EV2) or human CMG (Baris *et al*., 2022; Kang *et al*, 2012; Le *et al*, 2021) is very stable *in vitro* in the absence of DONSON, and reconstituted human replisomes containing CMG but lacking DONSON support a high rate of fork progression and DNA synthesis (Baris *et al*., 2022). Instead of preserving the integrity of CMG, our data suggest that DONSON is important for assembly of the CMG helicase in S-phase cells.

To explore this idea and determine which stage of CMG helicase assembly might require DONSON, we used Quantitative Image-Based Cytometry (Toledo *et al*, 2013) to monitor the association of MCM2-7, CDC45 and GINS with chromatin. Cells were given a pulse of EdU just before being analysed, to identify those cells that were undergoing DNA replication. When control cells were grown in the absence or presence of AGB1, the association of MCM2 with chromatin was maximal in G1 cells with a 2C DNA content and low EdU incorporation, and then gradually declined as cells progressed throughout S-phase (Fig 4A, cells shown in red), reaching a minimum in unreplicating G2-M cells with a 4C DNA content (Fig 4A, Control). A similar pattern of MCM2 chromatin binding was observed in *BromoTag-Donson* cells or *BromoTag-Psf1* cells (Fig EV5), even in the presence of AGB1 (Fig 4A, *BromoTag-*Psf1 and *BromoTag-Donson*). These data indicate that DONSON, like GINS, is dispensable for loading of the MCM2-7 helicase core onto chromatin at the end of mitosis.

**Figure 4.**
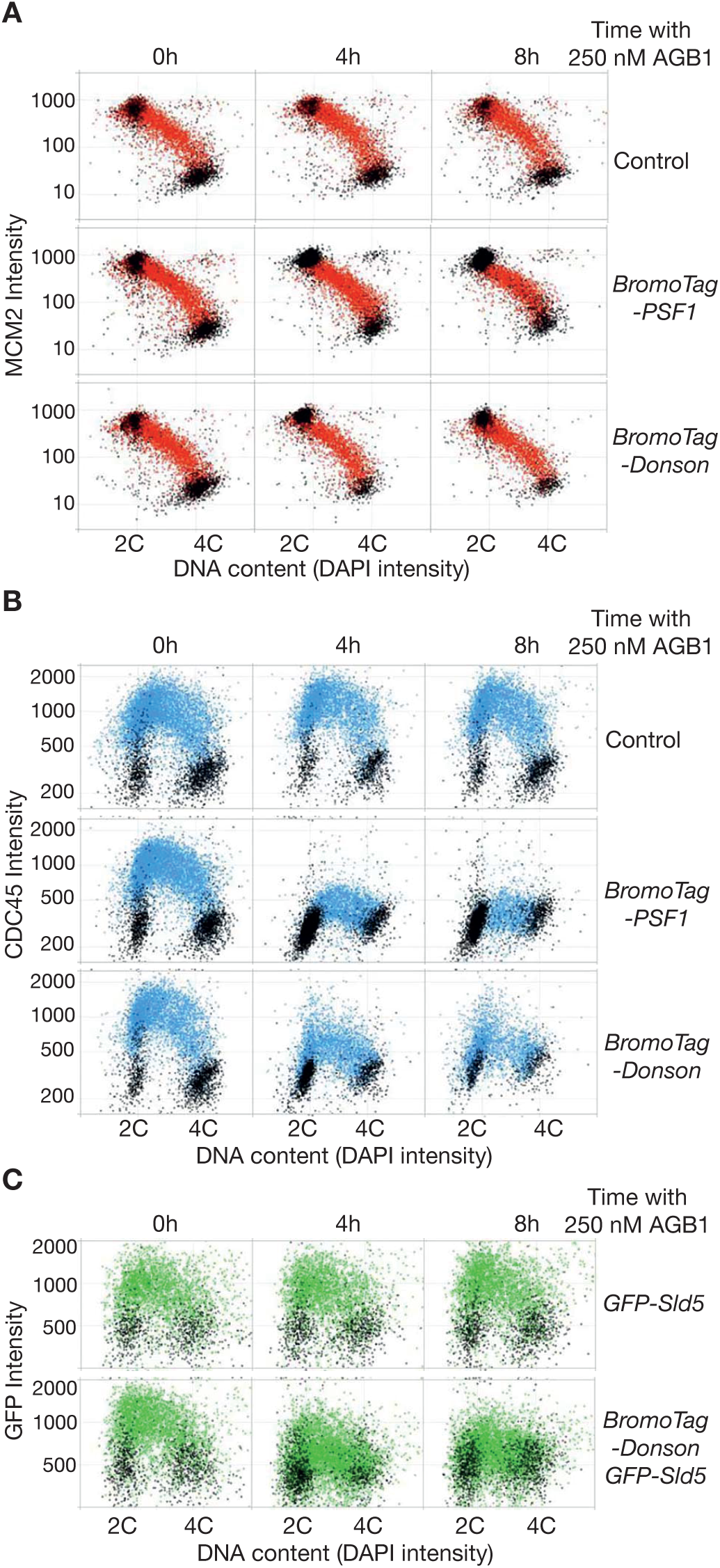
DONSON acts after loading of the MCM2-7 helicase core and is required for CMG helicase assembly. **(A)** The indicated cells were grown in the presence or absence of AGB1 as shown, before analysis of chromatin bound MCM2 by ‘Quantitative Imaging Based Cytometry’ (see Materials and Methods). Cells that incorporated EdU are shown in red, whereas cells that were not replicating are coloured black. **(B)** Similar analysis for CDC45 (cells that incorporated EdU are shown in blue with the remainder in black). **(C)** Equivalent experiment monitoring chromatin bound GFP-Sld5 (cells in green were EdU positive). Note that we were unable to monitor the chromatin association of GINS by immunofluorescence, perhaps due to masking of epitopes in the replisome, and thus utilised GFP-SLD5, as previously described (Villa *et al*., 2021).

Mouse ES cells have a short G1-phase, corresponding to cells that do not incorporate EdU and have little CDC45 on chromatin (Fig 4B, Control). Most cells are in S-phase with chromatin bound CDC45 (Fig 4B, cells shown in blue), before a short G2-phase and mitosis that once again correspond to minimal EdU incorporation and CDC45 not bound to chromatin. As expected, the association of CDC45 with chromatin was not affected when control cells were treated with AGB1 (Fig 4B, Control, 4-8h). In *BromoTag-Psf1* cells, CDC45 associated with chromatin when cells were grown in the absence of AGB1, with a very similar pattern to control cells (Fig 4B, *BromoTag-Psf1*, 0h). However, degradation of BromoTag-PSF1 blocked the association of CDC45 with chromatin (Fig 4B, *BromoTag-Psf1*, 4-8h), indicating that chromatin-bound CDC45 in this assay provides a readout for the CMG helicase at replication forks. Notably, degradation of BromoTag-DONSON also impaired the chromatin association of CDC45 (Fig 4B, *BromoTag-Donson*). Moreover, degradation of BromoTag-DONSON reduced the chromatin association of GINS in similar experiments (Fig 4C, cells shown in green - compare the *GFP-Sld5* control with *BromoTag-Donson GFP-Sld5* cells). Together with the immunoprecipitation experiments described above (Fig 3), these data indicate that DONSON is required for assembly of the CMG helicase on loaded MCM2-7 complexes, during DNA replication initiation in the mammalian cell cycle.

The importance of DONSON for CMG helicase assembly in mammalian cells provides a mechanistic explanation for why hypomorphic mutations in human *DONSON* lead to Meier Gorlin Syndrome, consistent with the fact that the other 12 Meier Gorlin genes encode components of the CMG helicase or factors that mediate or regulate helicase assembly (Nielsen-Dandoroff *et al*., 2023). Primordial Microcephalic Dwarfism syndromes such as Meier Gorlin result from defects in cell proliferation during embryogenesis, to which brain development is particularly sensitive (Klingseisen & Jackson, 2011). As noted above, *DONSON* mutations can also cause additional limb defects, as in Microcephaly with Short Stature and Limb Abnormalities (Schulz *et al*., 2018). Moreover, *DONSON* mutations that cause particularly severe defects in brain and limb development are associated with neonatal lethality, as in Microcephaly-Micromelia Syndrome (Evrony *et al*., 2017). Such conditions might involve *DONSON* mutations that produce particularly strong defects in CMG helicase assembly during embryogenesis or might be due to the disruption of additional functions of DONSON at replication forks.

A missense mutation in DONSON has also been linked to early-onset colorectal cancer (Tanskanen *et al*, 2015). In addition, multiple studies have found that over-expression of DONSON is associated with poor prognosis in a variety of human cancers (Klumper *et al*, 2020a; Klumper *et al*, 2020b; Qi *et al*, 2022; Yamada *et al*, 2020), as seen previously for other assembly factors of the CMG helicase (Hills & Diffley, 2014; Macheret & Halazonetis, 2015). Defining the molecular mechanism by which mammalian DONSON promotes CMG helicase assembly might inform the development of future anti-cancer therapies, based on synthetic lethality between partial inactivation of CMG assembly and the inherent replication defects that are retained by many cancer cells. DONSON function is likely to have been conserved during metazoan evolution, and new studies of DONSON orthologues in *Caenorhabditis elegans* (Xia *et al*, 2023) and *Xenopus laevis* (Hashimoto *et al*, 2023; Lim *et al*, 2023), reported during the preparation of this manuscript, indicate that DONSON recruits GINS to MCM2-7 during DNA replication initiation, using interfaces that are predicted by AlphaFold multimer (Evans *et al*, 2022) to be conserved in mammalian DONSON.

## Materials and Methods

Reagents and resources from this study are listed in Appendix Table S1.

### Culture of mouse ES cells

Mouse ES cells (E14tg2A) were maintained in Dulbecco’s Modified Eagle Medium (DMEM, ThermoFisher Scientific, 11960044) supplemented with 10% Foetal Bovine Serum (FBS, FCS-SA/500, LabTech), 5% KnockOut Serum Replacement (ThermoFisher Scientific, 10828028), 2 mM L-Glutamine (ThermoFisher Scientific, 25030081), 100 U/ml Penicillin-Streptomycin (ThermoFisher Scientific, 15140122), 1 mM Sodium Pyruvate (ThermoFisher Scientific, 11360070), a mixture of seven non-essential amino acids (ThermoFisher Scientific, 11140050), 0.05 mM β-mercaptoethanol (Sigma-Aldrich, M6250) and 0.1 μg/ml Leukemia Inhibitory Factor (MRC PPU Reagents and Services, DU1715). Cells were grown at 37°C in a humidified atmosphere of 5% CO_2_, 95% air. For passaging, cells were released from dishes using 0.05% Trypsin-EDTA (ThermoFisher Scientific, 25300054). Degradation of BromoTag-DONSON and BromoTag-PSF1 was induced by addition of 250 nM AGB1.

### Generation of plasmids expressing gRNAs for CRISPR-Cas9 genome editing

To facilitate genome editing by the *Streptococcus pyogenes* Cas9-D10A nickase, a pair of guide RNAs (gRNAs) was designed using ‘CRISPR Finder’ (https://wge.stemcell.sanger.ac.uk//find_crisprs), to target a specific site in the mouse genome. Each gRNA contains 20 nucleotides of homology, which in the genome is located immediately upstream of a 3 nucleotide ‘Protospacer Adjacent Motif’ (PAM) sequence that is required for cleavage by Cas9-D10A. Two annealed oligonucleotides containing the homology region were phosphorylated by T4 polynucleotide kinase (New England Biolabs, M201) and cloned into the vectors pX335 and pKN7 (see Appendix Table S1) after digestion by the Type IIS restriction enzyme BbsI as previously described (Pyzocha *et al*, 2014).

### Generation of donor vectors to introduce tags into mammalian genes by CRISPR-Cas9

To facilitate the introduction of purification tags, fluorescent proteins or degron cassettes into genomic loci in mammalian genomes, we developed a ‘mammalian toolkit’ for the modular construction of donor vectors *in vitro*. We adapted a previously described cloning system known as the ‘Yeast Toolkit’ (Lee *et al*, 2015), which employs Type IIS restriction enzymes that cut outside of their recognition sequence and so can be used to produce a range of DNA fragments with a bespoke set of cohesive ends, allowing the simple assembly of plasmids from multiple component parts. Vectors were generated to support the assembly of donor vectors that would allow the introduction of a range of tags at the amino or carboxyl terminus of the target protein. The resulting vectors are summarised in Appendix Figures S1-2, listed in Appendix Table S1, and are available from MRC PPU Reagents and Services (https://mrcppureagents.dundee.ac.uk/).

### Transfection of mouse ES cells and selection of clones

For each transfection, plasmids were generated (Appendix Table S1) that expressed the Cas9-D10A ‘nickase’, two gRNAs specific for the target locus and the Puromycin resistance gene. To introduce tags into genomic loci, an additional plasmid was generated that contained the sequence of the chosen tag with around 500-1000 bp homology to the target locus, together with the Hygromycin or Blasticidin resistance genes. Aliquots of 1.0 × 10^6^ cells were centrifuged at 350 × g for 5 minutes before resuspension in a mixture of 200 μl DMEM, 15 μl of 1 mg/ml linear Polyethylenimine (PEI; Polysciences, Inc, 24765-2) and 1-2 µg of each plasmid. The cells were then mixed gently by pipetting and incubated at room temperature for 30 minutes, before transfer to a single well of a 6-well plate precoated with 0.1% gelatin (Sigma, G1890) and containing 2 ml of complete DMEM medium. After incubation for 24 hours, cells were subjected to two 24-hour rounds of selection with fresh medium containing 2 μg/ml Puromycin (ThermoFisher Scientific, A1113802), followed by one to two weeks of selection with fresh medium containing 200-500 μg/ml Hygromycin (Invitrogen, ant-hg-5) or 7.5 μg/ml Blasticidin (Invitrogen, ant-bl-10p). The surviving cells were released from the well with 0.05% Trypsin-EDTA (ThermoFisher Scientific, 25300054), diluted from 10 to 10000 times with complete DMEM containing Hygromycin or Blasticidin, and plated on 10cm plates precoated with 0.1% gelatin. Subsequently, single colonies were picked and transferred to one well of a 6-well plate precoated with 0.1% gelatin. After 6 to 10 days, viable clones were expanded and monitored by immunoblotting, PCR and DNA sequencing of the target locus.

### PCR and sequence analysis for genotyping of mouse ES cells

Cells from a 10cm plate were released as described above. One third of the cells were pelleted and resuspended in 100 μl of 50 mM NaOH, before heating at 95°C for 15 minutes. The pH was then neutralised by addition of 11 μl of 1 M Tris-HCl (pH 6.9) and 0.5 µl used as a source of genomic DNA in 20 µl PCR reactions with PrimeStar Hot Start DNA polymerase (Takara Bio, R010A). Amplified DNA was subcloned with StrataClone Blunt PCR cloning kit (Agilent Technologies, 240207) and sequenced with M13 Forward, M13 Reverse primers, together with primers specific to the target locus.

### Synchronisation of mouse ES cells in mitosis

An aliquot of 2.5×10^6^ cells was seeded on a 10cm dish containing medium as described above. After 24 hours, cells were treated 1.25 mM Thymidine (Sigma-Aldrich, T9250) for 18 hours, to arrest in early S-phase. After 2 washes with Phosphate Buffered Saline (PBS), cells were released in fresh medium containing 5 μM STLC (Sigma-Aldrich, 164739) for 9 hours to arrest cells in mitosis. To degrade BromoTag-DONSON, 250 nM AGB1 (Tocris, 7686) was added for the last 2 hours of the STLC treatment. Cells were then washed 3 times with PBS and released in fresh medium, plus or minus 250 nM AGB1 as appropriate. A sample was taken every hour for 10 hours and processed for flow cytometry analysis.

### Flow cytometry

Cells from one 10 cm dish were released as described above and fixed with 1 ml 70% ethanol. For each time point, 100-300 µl of fixed cells were washed with PBS and resuspended in 1ml PBS containing 0.05 mg/ml propidium iodide (Sigma-Aldrich, P44170) and 0.05 mg/ml RNase A (ThermoFisher Scientific, EN0531). Cells were incubated at 24°C for 1 hour. The samples were then analysed with a FACSCanto II flow cytometer (Becton Dickinson) and the data processed with FlowJo software (https://www.flowjo.com/).

### Monitoring viability of mouse ES cells

An aliquot of 500 cells was seeded on a 10 cm dish. After 24 hours, AGB1 was added to the medium at a final concentration of 0, 25 or 250 nM. After 7 days at 37°C, the cells were washed with PBS and fixed with 20% methanol. Subsequently, cells were stained with 0.5% crystal violet solution (Sigma-Aldrich, HT90132) and the plates were scanned.

### Quantitative image-based cytometry (QIBC)

For the experiments in Figure 2C and Figure 4, mouse ES cells were seeded at 10,000 cells per well in clear-bottomed black 96-well plates (Greiner, 655090). Cells were grown for 24 h before addition of 250 nM AGB1 for 0, 4 or 8 hours, followed by treatment with 20 μM EdU for 20 minutes. To remove proteins not bound to chromatin, cells were pre-extracted for 5 minutes with cold CSK buffer (‘Cytoskeleton buffer’: 10 mM PIPES pH7, 100 mM NaCl, 300 mM sucrose, 3mM MgCl_2_) supplemented with 0.5% Triton X-100, 1mM CaCl_2_ and protease inhibitors (Merck, cOmplete EDTA-free, 1187358001; one tablet dissolved in 50 ml buffer), before fixation 20 minutes in 2% formaldehyde. For EdU detection, the far-red fluorescent dye Alexa-647 was covalently linked to EdU using the click-iT EdU Imaging Kit (Thermo Fisher, B10184). To detect CDC45, cells were incubated in methanol for 15 minutes at 4°C and then blocked with 5% BSA for 30 minutes, before incubation with a 1:200 dilution of a rabbit monoclonal anti-CDC45 antibody (Cell Signalling Technology, 11881) for 60 hours at 4°C, followed by 30 minutes at room temperature with donkey anti-rabbit IgG coupled to Alexa Fluor 488 (Invitrogen, A-21206), in the presence of 2.5 µg/ml DAPI (Thermo, 62248). For detection of MCM2, cells were processed as for CDC45 but without the methanol fixation step and with a 1:1000 dilution of mouse monoclonal anti-MCM2 (BD Biosciences, 610701) for 14 hours at 4°C, followed by 30 minutes at room temperature with donkey anti-mouse IgG coupled to Alexa Fluor 488 (Invitrogen, A-21202) together with 2.5 µg/ml DAPI (Thermo, 62248). GFP-SLD5 was detected using either of the above conditions used for CDC45 or MCM2. Finally, cells were washed and stored in PBS at 4°C until imaging by scanR High Content Screening Microscopy (Olympus). Images were analysed with the ScanR analysis software and data visualised in Tableau.

### Immunoprecipitations of protein complexes from extracts of mouse ES cells

Cells were seeded on 15cm plates at 1.5-2×10^7^ cells per dish. Typically, four dishes were used per sample (except Figure 1B where 16 plates were used). After 48 hours at 37°C, cells were released from the plates by incubation with PBS containing 1mM EDTA and 1mM EGTA for 10 minutes and harvested by centrifugation. Cells were then resuspended in one volume of lysis buffer (100 mM Hepes-KOH pH 7.9, 100 mM potassium acetate, 10 mM magnesium acetate, 2 mM EDTA, 10% glycerol, 0.1% Triton X-100, 2 mM sodium fluoride, 2 mM sodium β-glycerophosphate pentahydrate, 1 mM dithiothreitol, 1% Protease Inhibitor Cocktail (Sigma-Aldrich, P8215), 1x Complete Protease Inhibitor (Roche, 05056489001)) supplemented with 5 μM Propargyl-Ubiquitin (MRC PPU Reagents & Services, DU49003) to inhibit deubiquitylase activity. Chromosomal DNA was digested for 30 minutes at 4°C with 1600 U/ml of Pierce Universal Nuclease (ThermoFisher Scientific, 88702) before centrifugation at 20,000 × g for 30 minutes at 4°C. The resulting extract was added to magnetic beads (Life Technologies, Dynabeads M-270 Epoxy, 14302D) coupled to rabbit IgG (Sigma-Aldrich, S1265) as described previously (Xia *et al*, 2021) and then incubated for 2 hours at 4°C. Proteins were separated on a 4-12% Bis-Tris NuPAGE gel (Life Technologies, NP0301) in 1X MOPS and detected by immunoblotting.

For mass spectrometry analysis, 16 dishes were used per sample. Gels were stained with colloidal Coomassie (ThermoFisher Scientific, Simply Blue, LC6060) and each sample lane was then cut into 40 bands before digestion with trypsin. Peptides were analysed by nano liquid chromatography tandem mass spectrometry with an Orbotrap Velos (MS Bioworks, https://www.msbioworks.com/).

### Expression and purification of recombinant mouse DONSON in bacteria

The coding sequence of mouse *Donson* was amplified from XpressRef Universal Total mouse RNA (QIAGEN, 338114) by RT-PCR (Takara, RR014) and cloned by Gibson assembly into the vector pK27SUMO. The resulting plasmid was transformed into Rosetta (DE3) pLys (Novagen, 70956) and a 100 ml overnight culture was grown overnight at 37°C in LB medium supplemented with kanamycin (50 ug/ml) and chloramphenicol (35 ug/ml). The next day, the culture was diluted 20 times in 1L LB medium with kanamycin and chloramphenicol until the OD 600 reached 0.8-1.0. Expression of mouse DONSON was then induced by addition of 1 mM Isopropyl β-d-1-thiogalactopyranoside (IPTG) for 3 hours at 37°C. Cells were then centrifugated for 10 minutes in a JLA-9.1000 rotor (Avanti J-25 centrifuge, Beckman) at 5180 × g and the pellet was stored at -20°C.

Subsequently, the frozen cell pellet was thawed and resuspended in 25 ml Lysis buffer (50 mM Tris-HCl pH 8, 500 mM NaCl, 20 mM imidazole, 10% glycerol, 0.02% NP-40, 0.5 mM TCEP, 1x Complete Protease Inhibitor (Roche, 05056489001)) supplemented with 1 mg/ml Lysozyme (Sigma-Aldrich, 000000010837059001) and incubated on ice for 30 minutes. The extract was then sonicated for 2 minutes (5 seconds on, 5 seconds off) at 40% on a Branson Digital Sonifier and clarified by centrifugation for 30 minutes at 27,216 × g at 4°C in an SS-34 rotor (Avanti J-25 centrifuge, Beckman).

The supernatant was added to 1.5 ml of Ni-NTA resin (Qiagen, 30210). After 2 hours incubation on a rotating wheel at 4°C, the beads were washed extensively with a total of 80 column volumes of Lysis buffer (protease inhibitors were omitted in the last wash). Subsequently, His-SUMO-DONSON was eluted with 5 column volumes of elution buffer (50 mM Tris-HCl pH 8, 500 mM NaCl, 400 mM imidazole, 10% glycerol, 0.02% NP-40, 0.5 mM TCEP). The elution fractions were pooled and the His-SUMO tag was cleaved with 400 μg of ULP1 SUMO protease (MRC PPU Reagents & Services, DU39129) overnight at 4°C. The sample was then concentrated with an Amicon Ultra-15 Centrifugal Filter Unit with a 30 kDa cut-off (Millipore, UFC903024) and loaded onto a 24 ml Superdex 200 column equilibrated in gel filtration buffer (50 mM Tris-HCl pH 8, 500 mM NaCl, 10% glycerol, 0.02% NP-40, 0.5 mM TCEP). Fractions containing purified DONSON were then pooled, aliquoted and stored at -80°C.

### Expression of mouse CMG in budding yeast cells

The coding sequence of the 11 subunits of mouse CMG were codon optimised for high-level expression in budding yeast cells and produced by DNA synthesis (GenScript Biotech). A previously described ‘yeast toolkit’ (Lee *et al*., 2015), based on Type IIS restriction enzymes that allow the seamless joining of multiple DNA fragments *in vitro*, was used to generate two plasmids that each have three transcription units under control of the *GAL1,10* bi-directional promoter (Appendix Table S1, pCE297 and pCE1259). The first plasmid supported expression of the six MCM2-7 proteins (pCE207) and was integrated at the *leu2* locus in the haploid yeast strain YCE923, producing the strain YCE1206. The second plasmid supported expression of CDC45 and the four GINS subunits (pCE297) and was integrated at the *HO* locus of YCE1206, using CRISPR-Cas9 as described previously. The resulting strain was crossed to YJF1 to produce the final strain YCE1259.

### Purification of recombinant mouse CMG

A 15L culture of the budding yeast strain YCE1259 was grown at 30°C in rich medium (1% Yeast Extract, 2% Peptone) supplemented with 2% Raffinose. When cell density reached ∼2.5 × 10^7^ cells/ml, 2% Galactose was added to induce expression of the CMG subunits for 6 hours at 30°C. Cells were then harvested by centrifugation at 5180 × g for 10 minutes at 4°C. Cells were washed with CMG buffer/300 mM KCl (25 mM Hepes-KOH pH 7.6, 10% glycerol, 2 mM MgOAc, 0.02% Tween-20, 1mM DTT) and the pellet resuspended with 0.3 volumes of CMG buffer/300 mM KCl supplemented with 1x Complete Protease Inhibitor (Roche, 05056489001). The cell suspension was then frozen dropwise in liquid nitrogen. The frozen yeast cells were ground with a freezer mill (SPEX CertiPrep 6850 Freezer Mill) using 4 cycles of 2 minutes at a rate of 15 cycles per second. The thawed yeast cell powder was resuspended in one volume of CMG buffer/300 mM KCl supplemented with protease inhibitor and the extract clarified by centrifugation in 1 Ti 45 rotor at 235, 000 × g at 4°C for 1 hour (Optima L-90K Ultracentrifuge). The supernatant was mixed with 3 ml of IgG Sepharose Fast Flow beads (GE Healthcare, 17096901) equilibrated with CMG buffer/300 mM KCl and incubated for 2 hours at 4°C on a rotating wheel. After extensive washes, firstly with 35 column volumes of CMG buffer/300 mM KCl containing protease inhibitor and then with 35 column volumes of CMG buffer/200 mM KCl, the beads were resuspended in one volume CMG buffer/200 mM KCl supplemented with 192 μg of TEV protease and the mixture was incubated overnight at 4°C. The eluate was then loaded onto a 1 ml HiTrap Q HP (Sigma-Aldrich, GE17-1153-01) equilibrated in CMG buffer/200 mM KCl. Purified CMG was then eluted with a gradient from 200 to 600 mM KCl over 20 column volumes. Fractions containing all 11 subunits were pooled and concentrated with an Amicon Ultra-15 Centrifugal Filter Unit with a 100 kDa cut-off (Millipore, UFC910024) and then loaded onto a 24 ml Superose 6 column equilibrated in CMG buffer/200 mM KCl. Fractions containing all 11 subunits were again pooled and concentrated with an Amicon Ultra-15 Centrifugal Filter Unit 100 kDa cut-off (Millipore, UFC910024) and subsequently dialysed against CMG buffer/250 mM KOAc overnight at 4°C. The complex was then aliquoted, snap-frozen and stored at -80°C.

### *In vitro* binding assay for mouse CMG and DONSON

For the experiment in Figure 1D, the indicated amounts of purified recombinant mouse DONSON and CMG were incubated in 25 mM Hepes-KOH pH7.6, 200 mM KOAC, 0.02% NP-40, 10 mM MgOAc, 1 mM DTT and 0.1 mg/ml BSA for 30 minutes on ice. Subsequently, 5 μl of magnetic beads (Life Technologies, Dynabeads M-270 Epoxy, 14302D) coupled to mouse MCM6 antibody was added to the proteins and incubated for 1 hour on ice with occasional mixing. After 2 washes with the above buffer, the bounds proteins were eluted from the beads with 15 μl of 1x Laemmli buffer by heating at 95°C for 5 minutes. Proteins were resolved in a 4-12% Bis-Tris NuPAGE gel (Life Technologies, NP0301) in 1X MOPS and detected by immunoblotting.

### SDS-PAGE and immunoblotting

Proteins were resolved in 4-12% Bis-Tris NuPAGE gels (Life Technologies, NP0301) in 1X MOPS buffer and then transferred onto a nitrocellulose membrane (ThermoFisher Scientific, IB301031) using the iBlot Dry transfer System (ThermoFisher Scientific). Membranes were blocked with 5% dried milk in TBST buffer (20 mM Tris base, 137 mM Sodium chloride pH7.6, 0.1% Tween 20) for 1 hour at room temperature. Primary antibodies were diluted at the concentrations listed in Appendix Table S1 with 5% dried milk in TBST and then incubated with membranes overnight at 4°C. The membranes were then washed three times for 10 minutes with TBST, before incubation with secondary antibody at the concentrations listed in Appendix Table S1 with 5% dried milk in TBST, for 1 hour at room temperature. The membranes were washed again three times for 10 minutes each with TBST and proteins were detected on the Azure 300 imaging system (Cambridge Bioscience) using ECL Western Blotting Detection Reagent (VWR, RPN2106).

### Live imaging of DONSON-GFP and mCherry-PSF1 by spinning disk confocal microscopy

For the experiment in Fig 1D, mouse ES cells expressing DONSON-GFP and mCherry-PSF1 from the endogenous loci were grown on ‘µ-Slide 4-well’ (Ibidi, 80426) with ‘no phenol red DMEM medium’ (ThermoFisher Scientific, 21063029) supplemented as described above. For p97 inhibition, cells were treated with 5 µM CB-5083 for 3 hours before imaging.

Confocal images of live cells were acquired with a Zeiss Cell Observer SD microscope with a Yokogawa CSU-X1 spinning disk, using a HAMAMATSU C13440 camera with a PECON incubator, a 60X 1.4-NA Plan-Apochromat oil immersion objective, and excitation and emission filter sets for GFP and mCherry. Images of live mouse ES cells were acquired using ‘ZEN blue’ software (Zeiss) and processed with ImageJ software (National Institutes of Health) as previously described (Sonneville *et al*., 2017).

## Data availability

This study includes no data deposited in external repositories.

## Acknowledgements

We thank Fabrizio Villa for help with growth and manipulation of mouse ES cells in the early stages of this work, Tobias Meyer’s group for help with conditions for CDC45 immunofluorescence and MRC PPU Reagents and Services (https://mrcppureagents.dundee.ac.uk) for antibody production. We are grateful for the support of the Medical Research Council (core grant MC_UU_12016/13 to KL), Cancer Research UK (Programme Grants C578/A24558 to KL and CDA-21782 to CA), the European Research Council (ERC-stg-715127 to CA) and the Japan Society for the Promotion of Science (JSPS Overseas Research Fellowship 202160572 to RF). Materials generated in this study are listed in Appendix Table S1 and are available from MRC PPU Reagents and Services (https://mrcppureagents.dundee.ac.uk) or upon request.

---

**Figure EV1.**
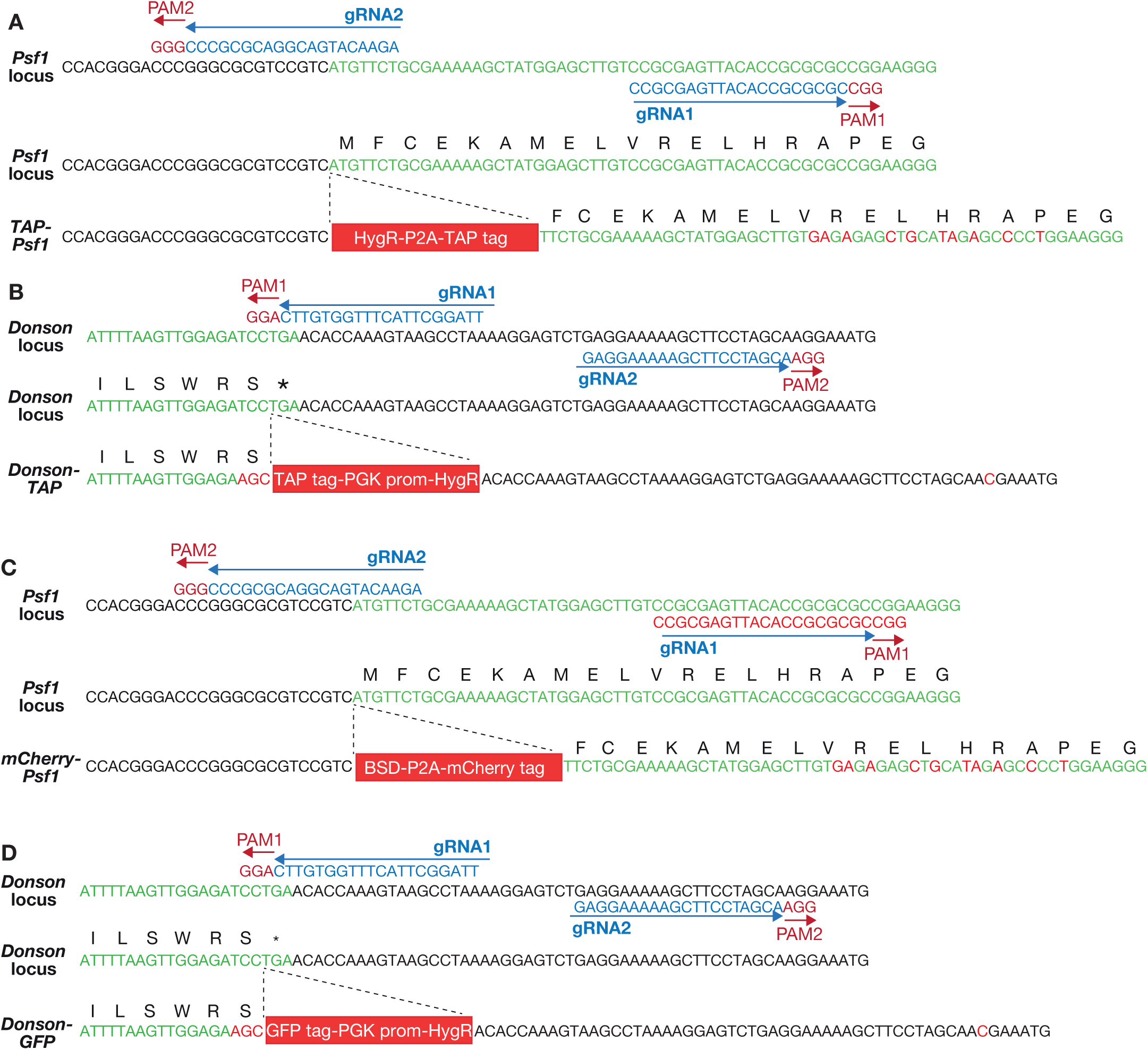
Genome editing of the *Psf1* and *Donson* loci in mouse embryonic stem cells by CRISPR-Cas9-D10A. **(A)** N-terminal tagging of PSF1 with Tandem Affinity Purification tag (TAP tag). Plasmids expressing *Psf1* gRNA1 (pCE319) and *Psf1* gRNA2 (pCE320), together with the Cas9-D10A nickase and a donor vector (pCE318), were used to introduce the indicated cassette before the initiator ATG of the *Psf1* gene. The tag comprises a Hygromycin resistance marker followed by the P2A sequence (that causes ribosomal skipping during translation) and the TAP tag. In this way, the cells express TAP-PSF1 and the Hygromycin resistance marker as separate proteins. **(B)** C-terminal tagging of DONSON with TAP. Plasmids expressing *Donson* gRNA1 (pCE308) and *Donson* gRNA2 (pCE308), together with the Cas9-D10A nickase and the donor vector pCE315, were used to introduce the indicated cassette in place of the STOP codon at the 3’ end of the *Donson* gene. The inserted cassette comprises the TAP tag (in frame with DONSON) followed by Hygromycin resistance marker under control of the *PGK1* promotor. **(C)** N-terminal tagging of PSF1 with mCherry. As for (A) above, except that the tag comprises the Blastocidin S deaminase gene from *Aspergillus terreus*, followed by the P2A sequence and the mCherry fluorescent protein (donor vector pCE334). In this way, the cells express mCherry-PSF1 and the Blastocidin resistance marker as separate proteins. (**D**) C-terminal tagging of DONSON with GFP. As for (B) above but with GFP in place of TAP (donor vector pCE333).

**Figure EV2.**
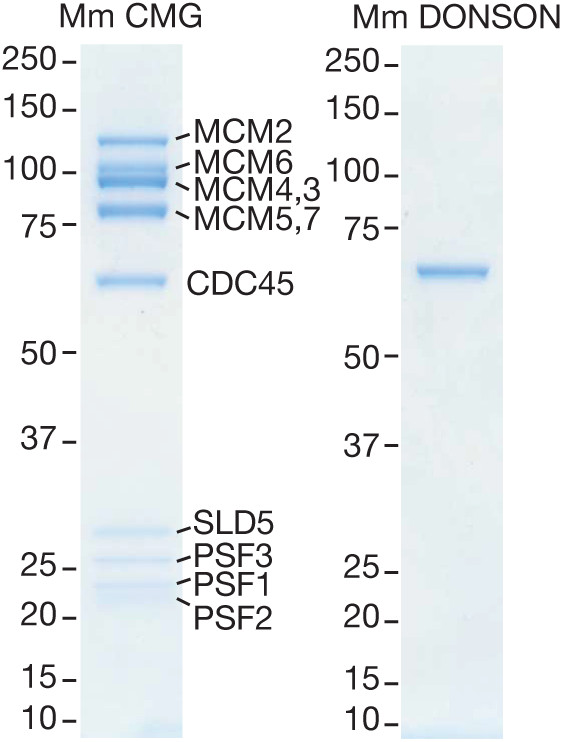
Purified recombinant versions of mouse CMG and DONSON. Mouse CMG was expressed and purified from budding yeast cells, whereas DONSON was expressed and purified from bacteria, as described in Materials and Methods. The purified proteins were resolved by SDS-PAGE and stained with colloidal Coomassie blue.

**Figure EV3.**
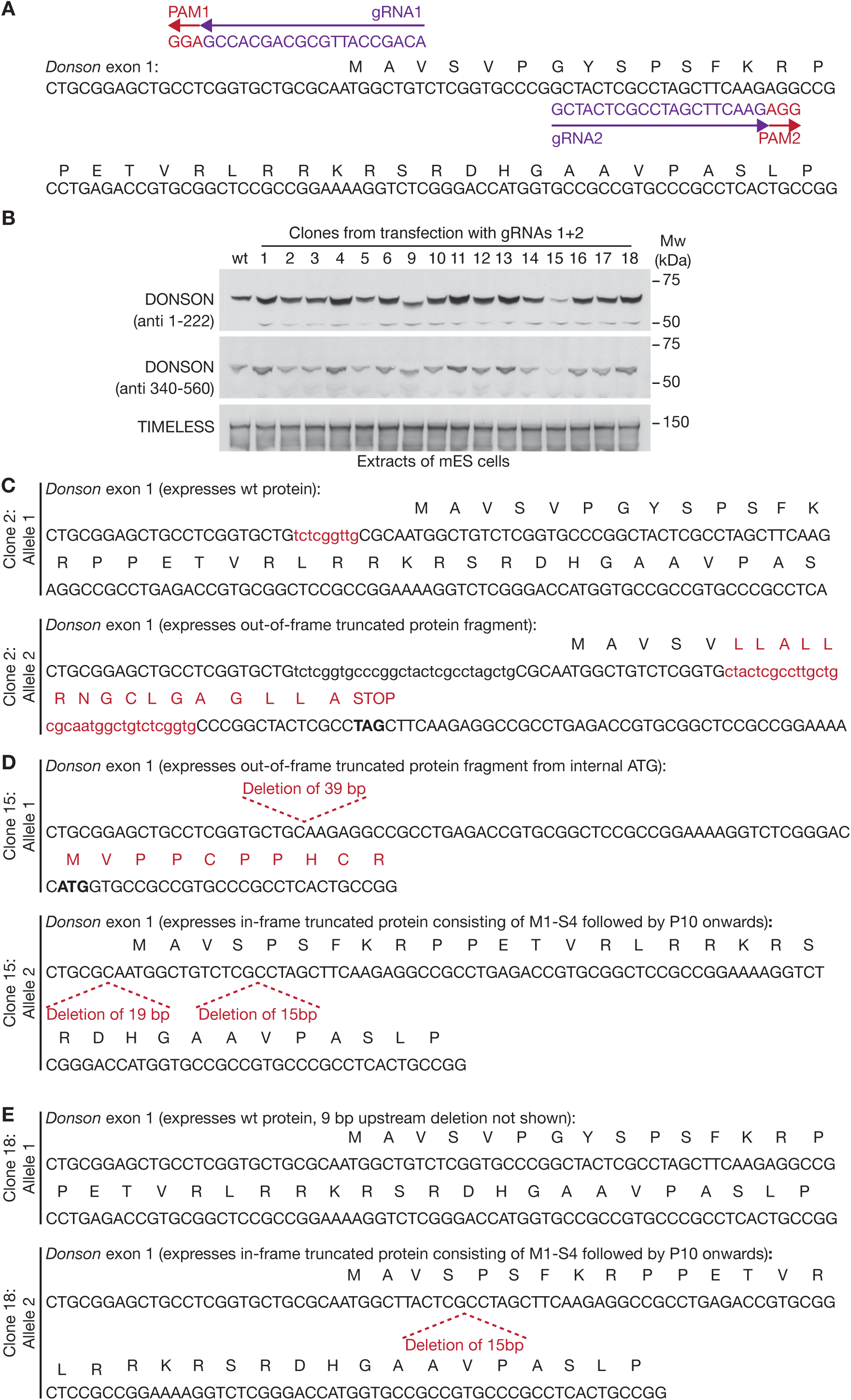
Analysis of viable clones with small deletions at the beginning of the *Donson* coding sequence in mouse ES cells. **(A)** Location of Donson gRNA1 (pJA5) and Donson gRNA2 (pJA6) are shown, together with the associated Protospacer Adjacent Motifs in the genomic sequence. Cas9-D10A cuts close to the PAM sequences and thus drives the formation of small deletions or insertions when the cut sequences are repaired. **(B)** Immunoblot analysis of selected clones, after transfection of mouse ES cells with *Donson* gRNAs 1+2. (**C**-**E**) DNA sequence analysis of both alleles of *Donson* in selected clones from (B).

**Figure EV4.**
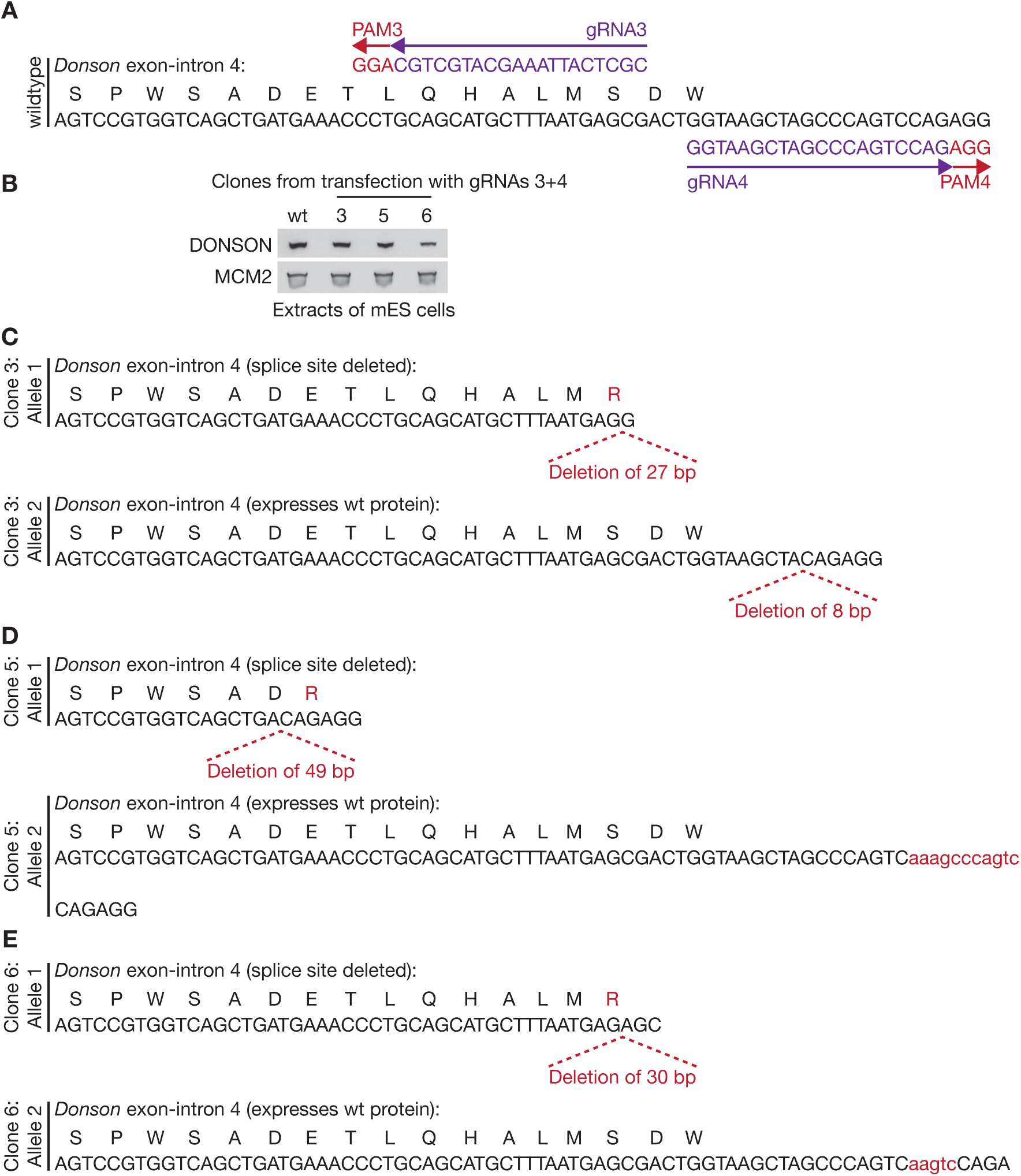
Analysis of viable clones with small deletions at the end of exon 4 of the *Donson* gene in mouse ES cells. **(A)** Location of Donson gRNA3 (pJA11) and Donson gRNA4 (pJA12) are shown, together with the associated Protospacer Adjacent Motifs in the genomic sequence. Cas9-D10A cuts close to the PAM sequences and thus drives the formation of small deletions or insertions when the cut sequences are repaired. **(B)** Immunoblot analysis of selected clones, after transfection of mouse ES cells with *Donson* gRNAs 3+4. (**C**-**E**) DNA sequence analysis of both alleles of *Donson* in clones from (B).

**Figure EV5.**
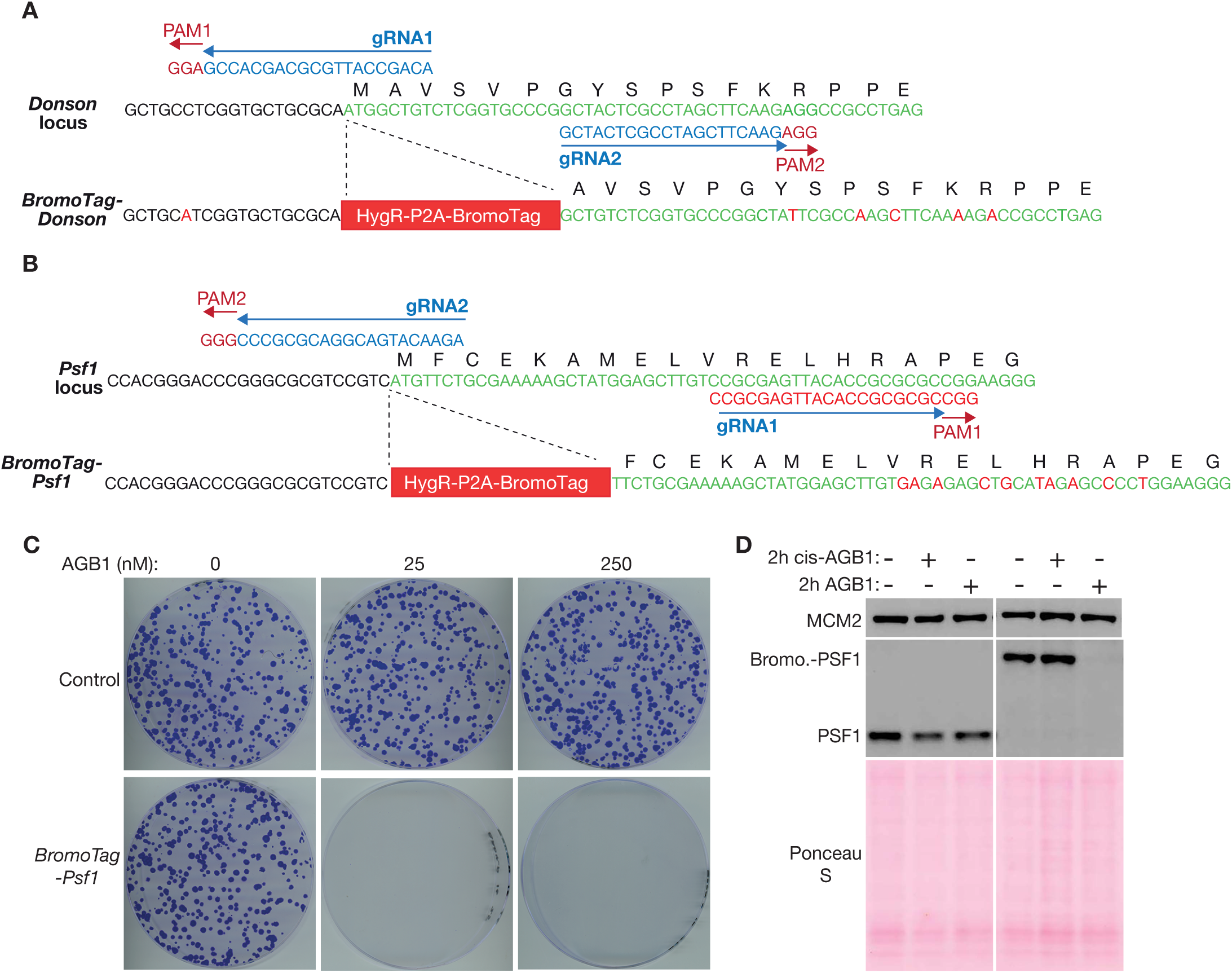
Genome editing of mouse embryonic stem cells to introduce the BromoTag degron cassette into the Donson and Psf1 loci. **(A)** N-terminal tagging of DONSON with the BromoTag degron cassette. Plasmids expressing *Donson* gRNA1 (pJA5) and *Donson* gRNA2 (pJA6), together with the Cas9-D10A nickase, were used to introduce the indicated cassette before the initiator ATG of the *Donson* gene. The tag comprises a Hygromycin resistance marker followed by the P2A sequence (that causes ribosomal skipping during translation) and the BromoTag. In this way, *BromoTag-Donson* cells express BromoTag-DONSON and the Hygromycin resistance marker as separate proteins. **(B)** N-terminal tagging of PSF1 with the BromoTag degron cassette. Plasmids expressing *Psf1* gRNA1 (pCE319) and *Psf1* gRNA2 (pCE320), together with the Cas9-D10A nickase, were used to introduce the indicated cassette before the initiator ATG of the *Psf1* gene. *BromoTag-Psf1* cells express BromoTag-PSF1 and the Hygromycin resistance marker as separate proteins. **(C)** Control and *BromoTag-Psf1* cells were grown for 7 days in the presence of the indicated concentrations of AGB1, before staining with crystal violet solution as described in Materials and Methods. **(D)** Immunoblots showing the degradation of BromoTag-PSF1 after treatment with 250 nM AGB1 for 2 hours. cis-AGB1 provides a negative control and has the *cis*-instead of *trans*-hydroxyproline group, abrogating binding to the VHL component of the ubiquitin ligase CUL2^VHL^.

## Appendix

**Appendix Figure S1.**
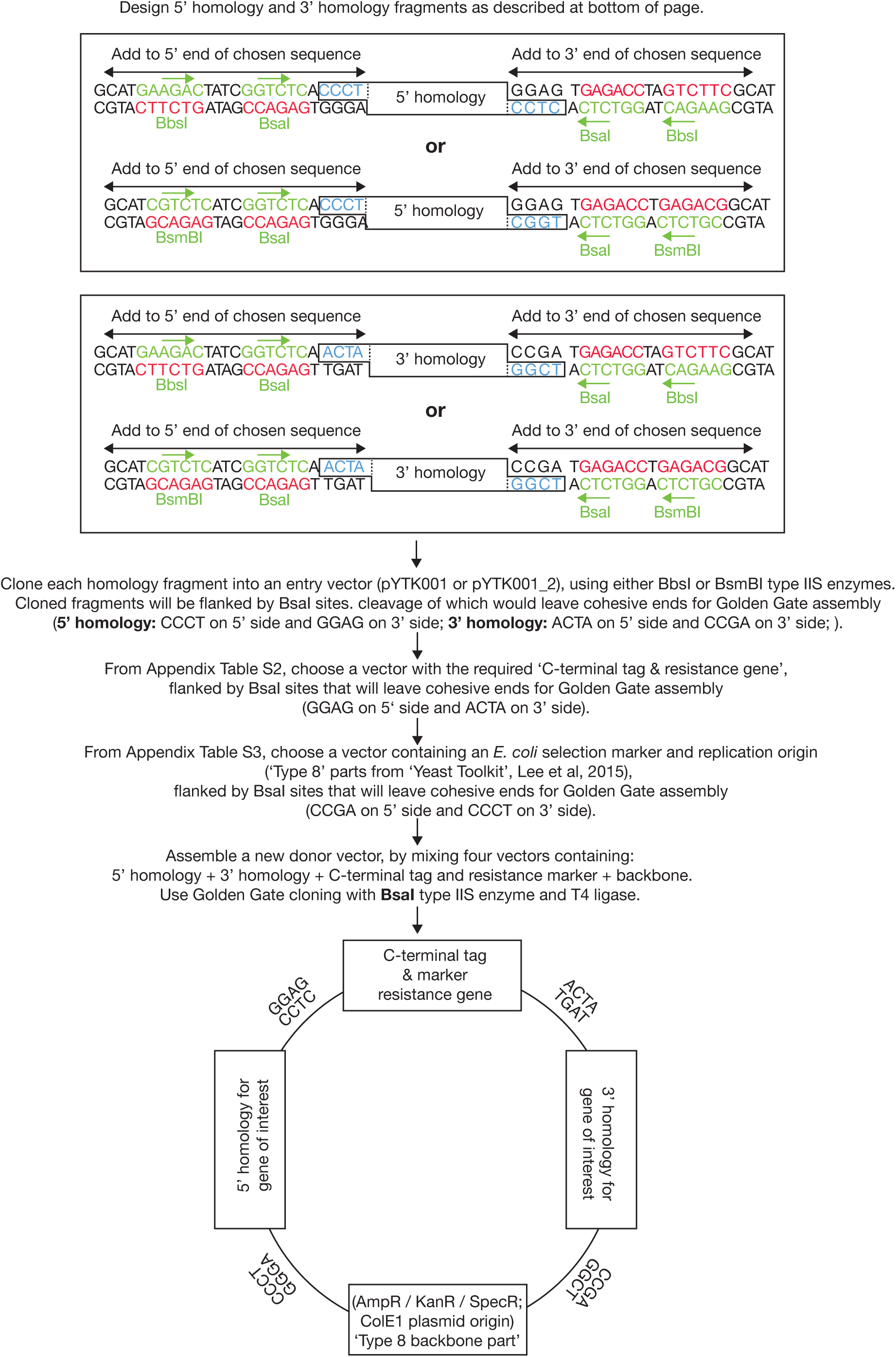
Design of 5’ and 3’ homology fragments for assembly of donor vectors for C-terminal tagging in mammalian cells.

**Appendix Figure S2.**
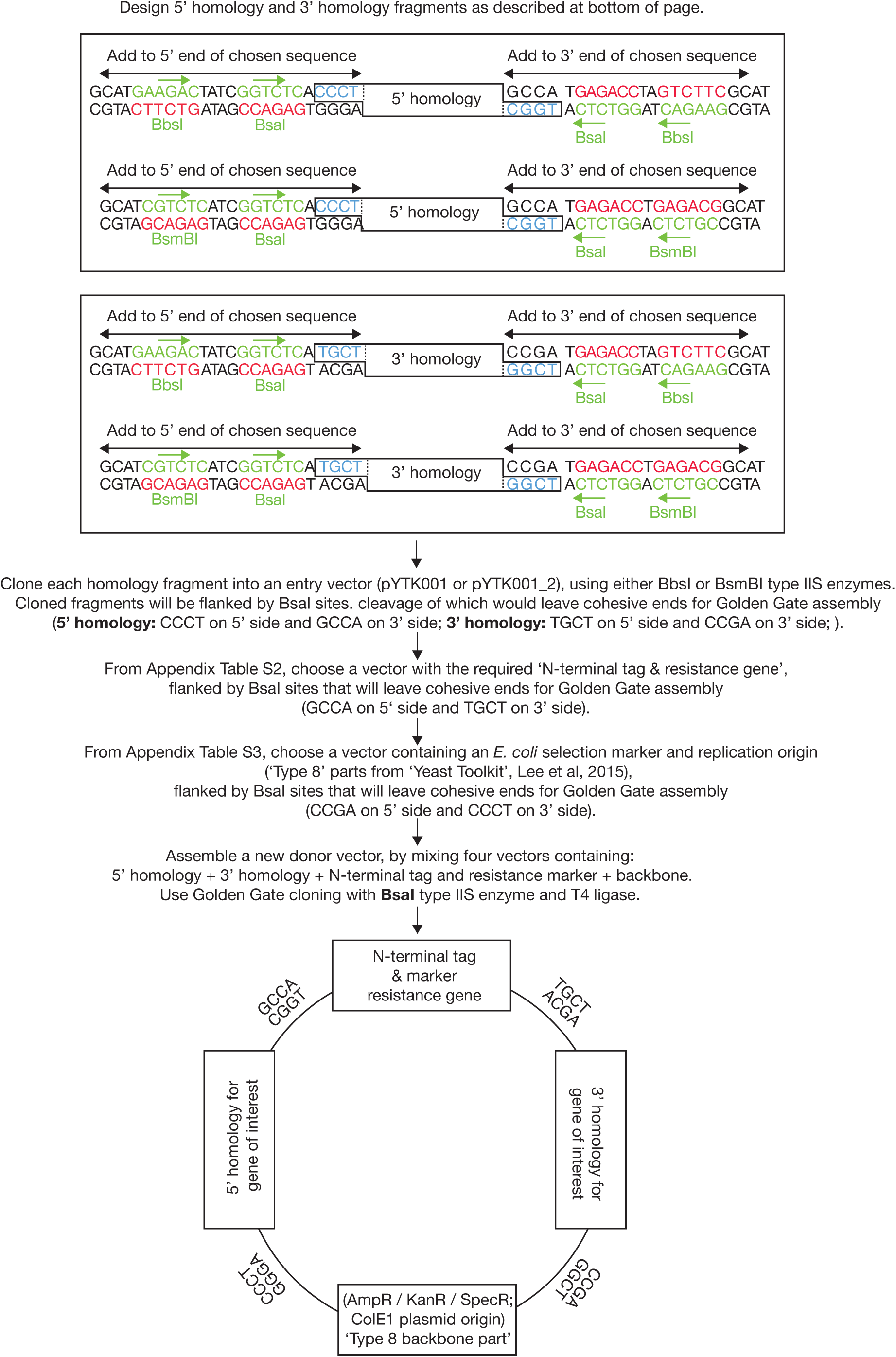
Design of 5’ and 3’ homology fragments for assembly of donor vectors for N-terminal tagging in mammalian cells.

**Appendix Table S1.**
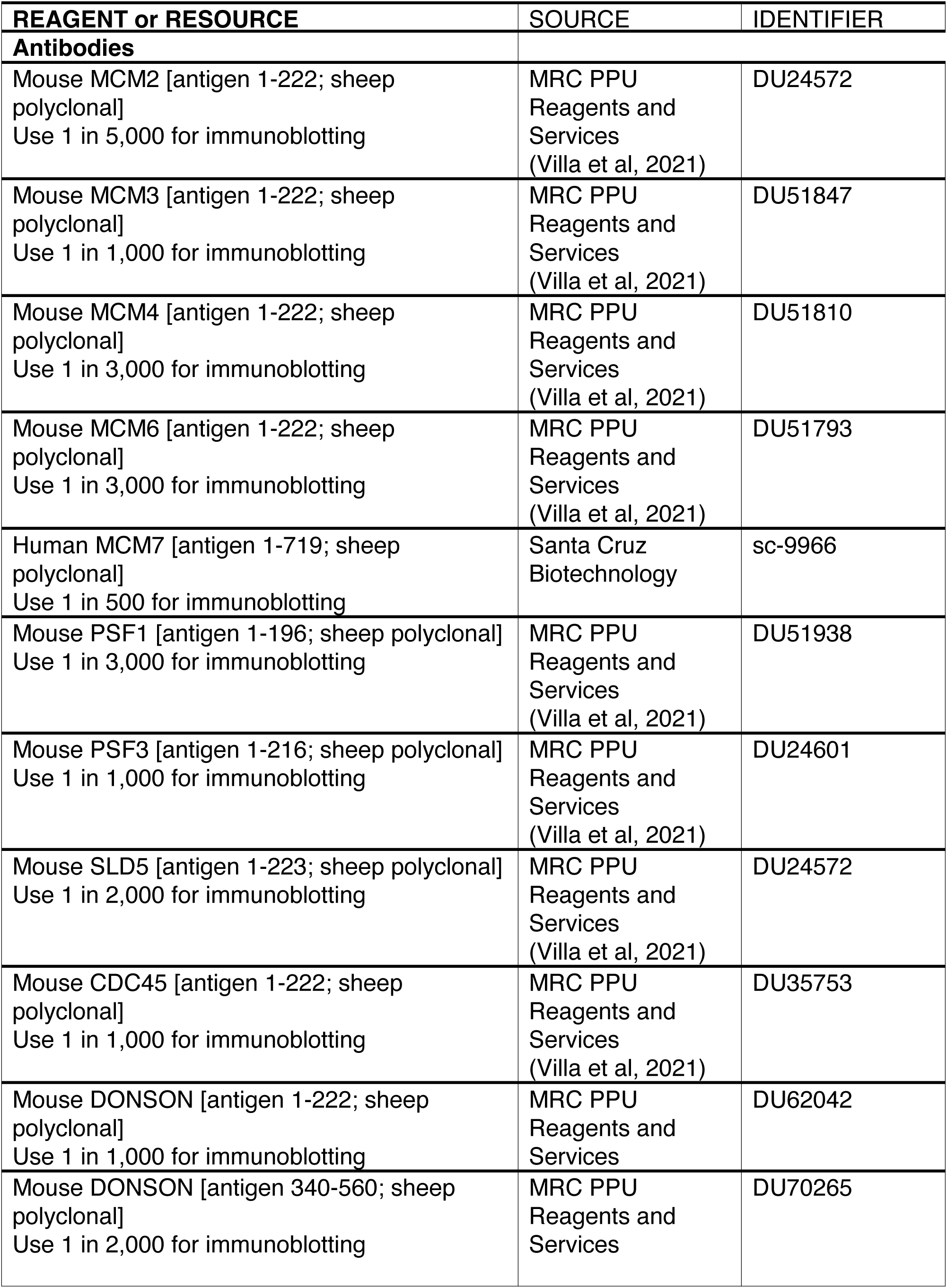
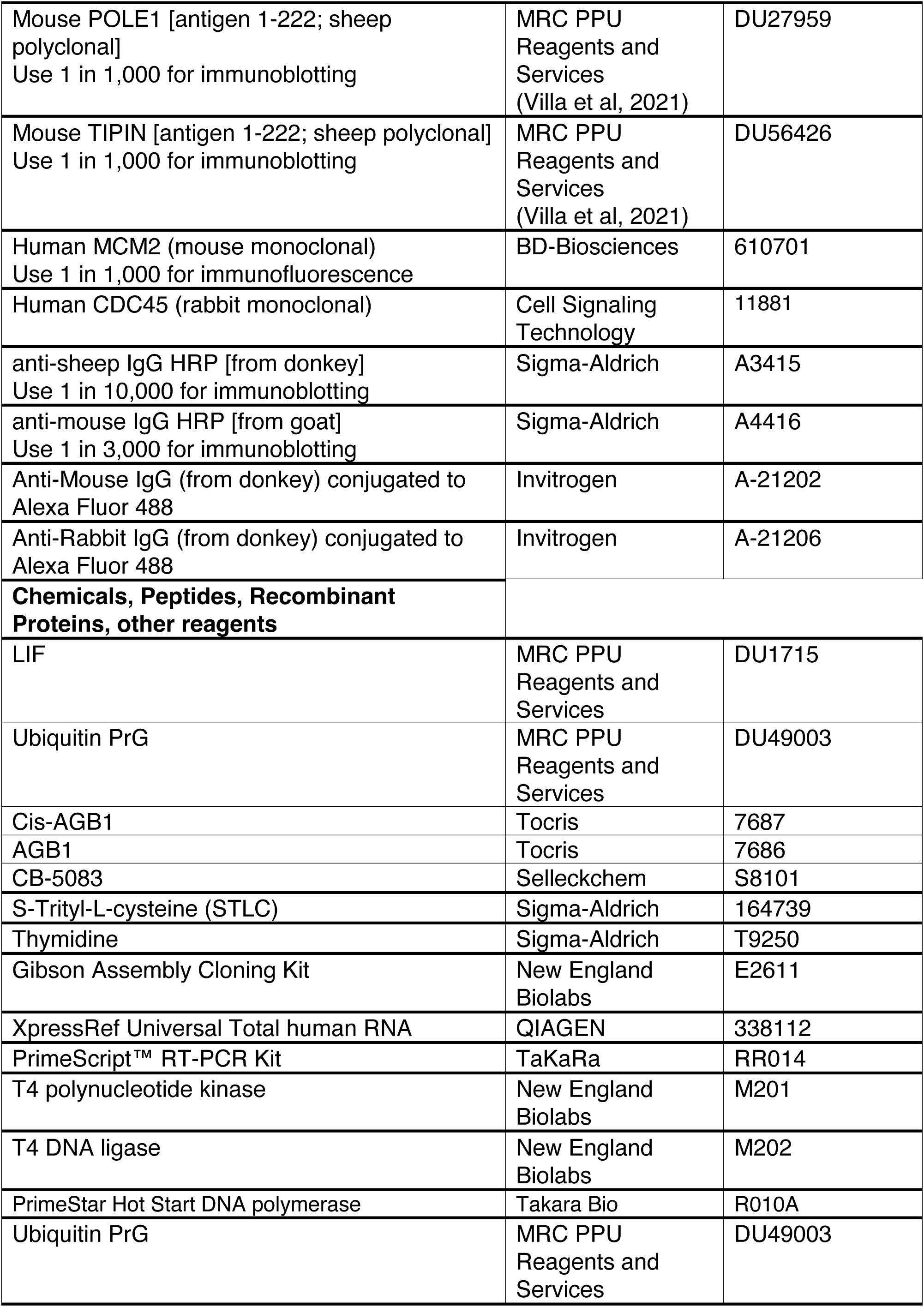
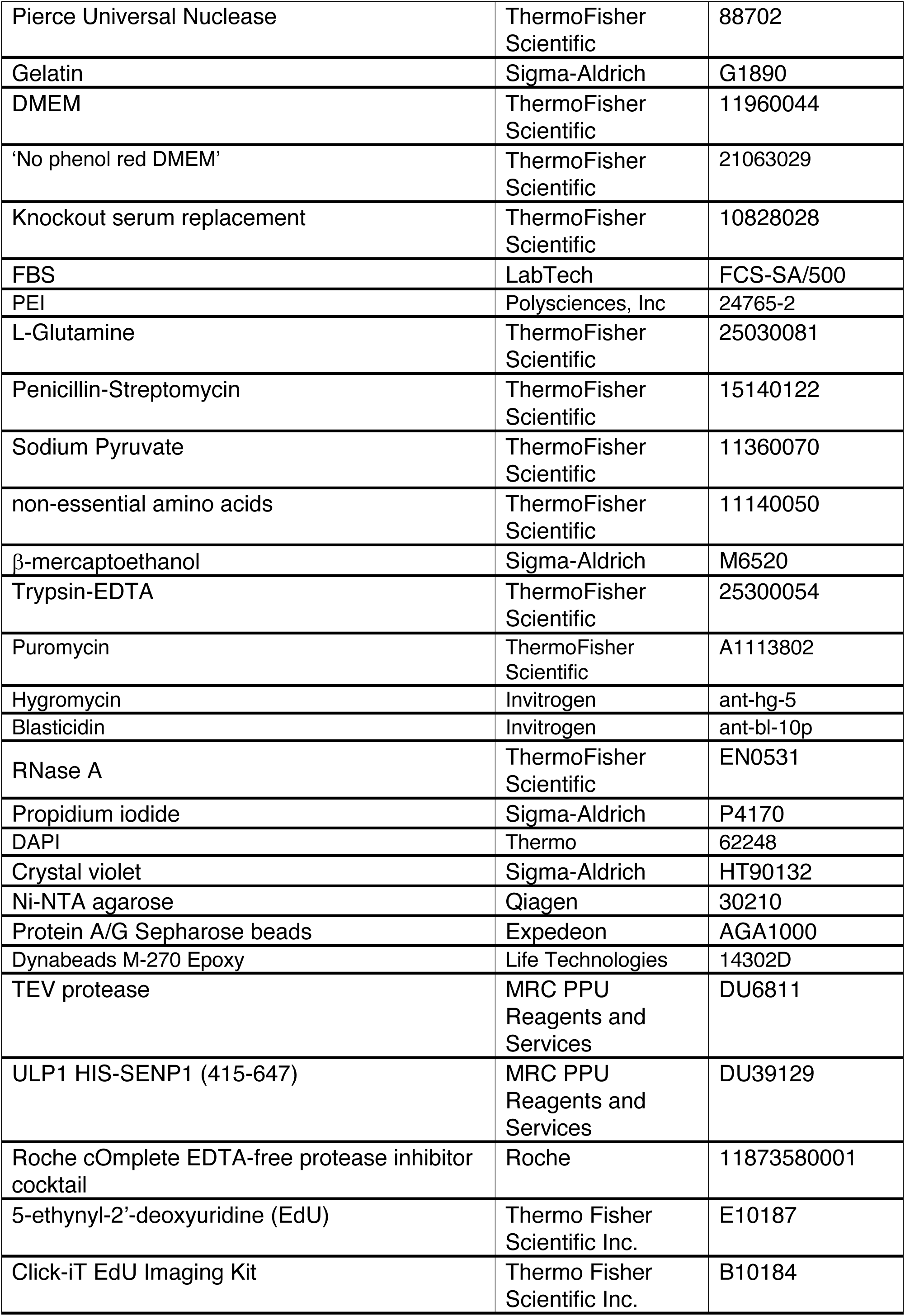
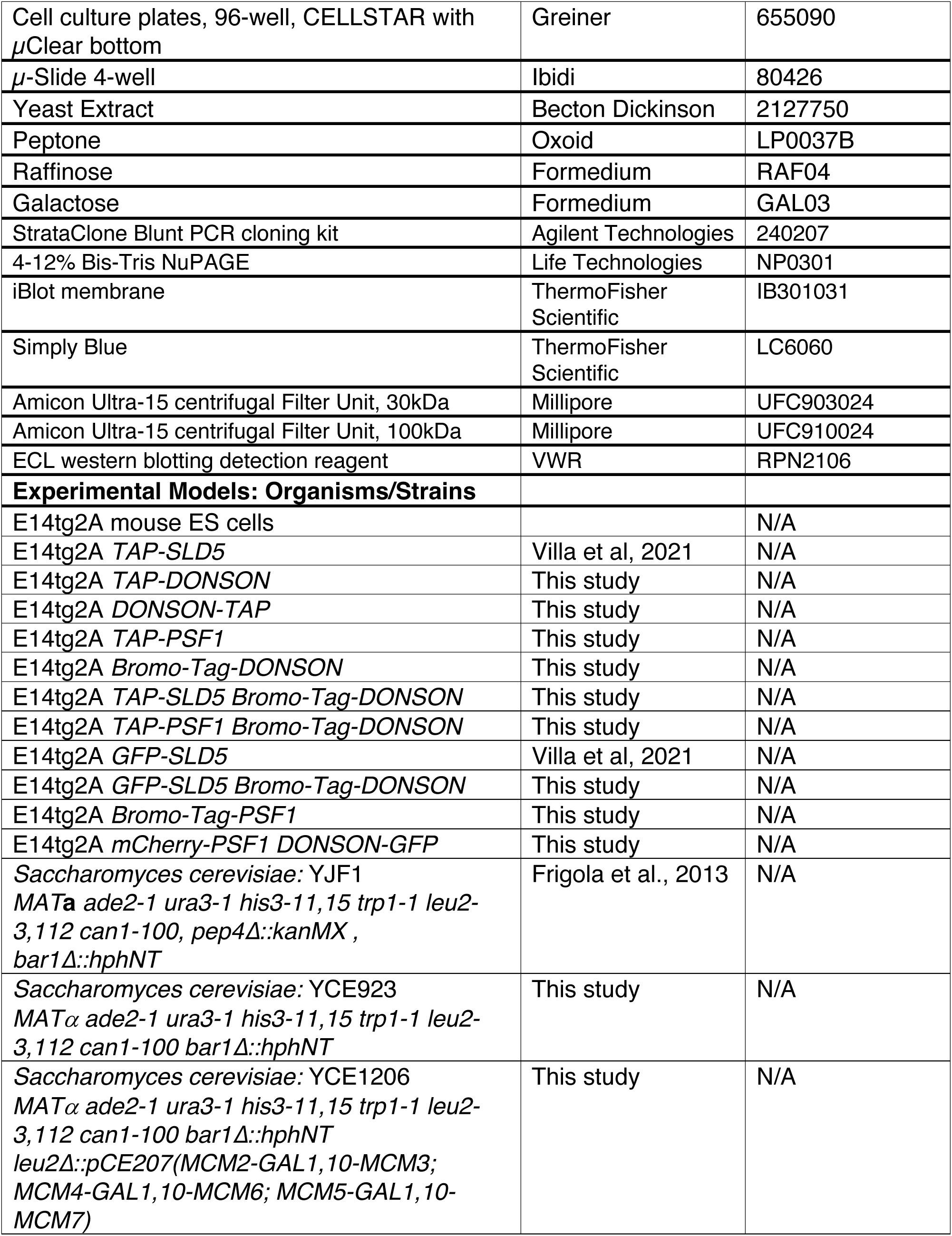
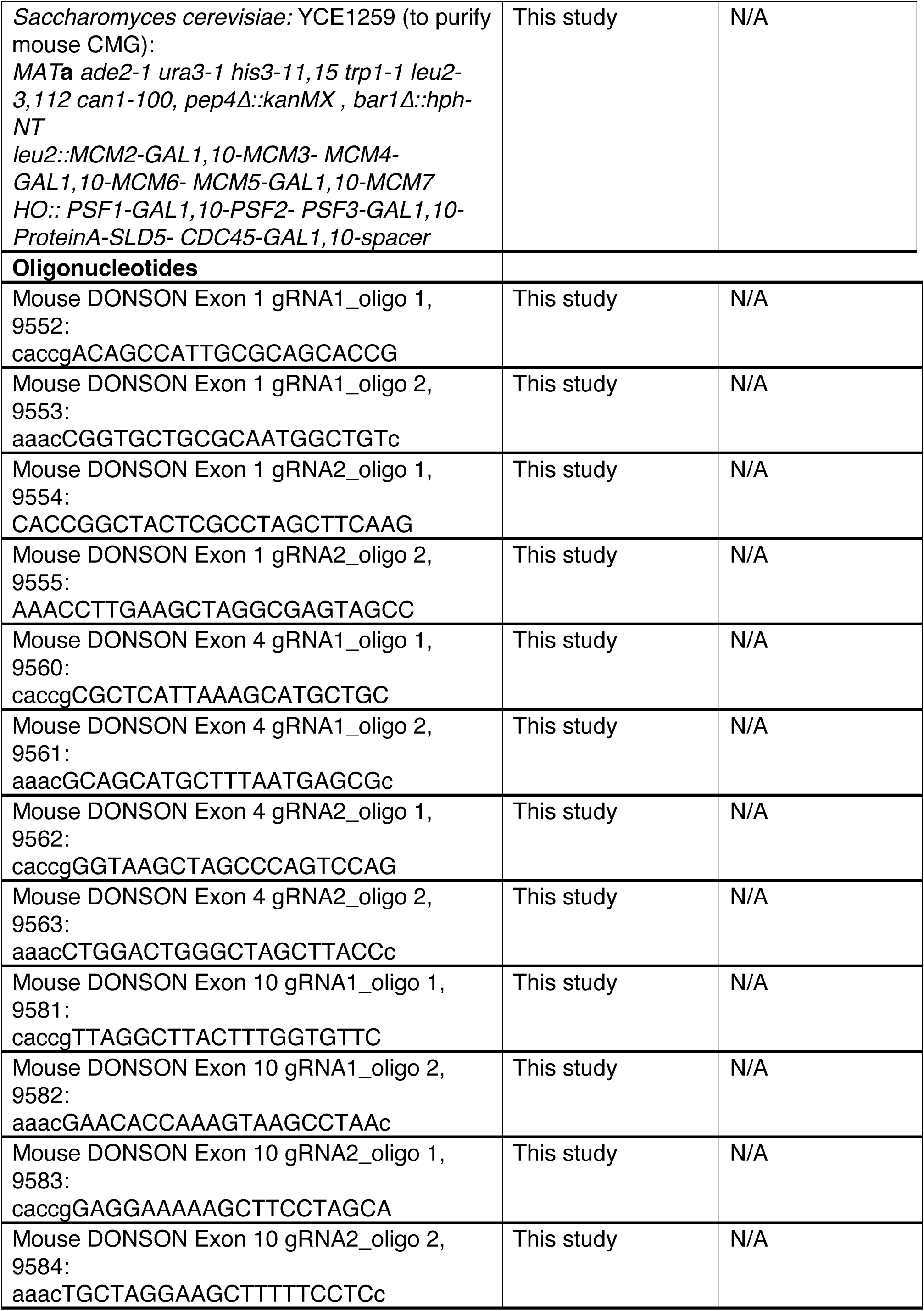
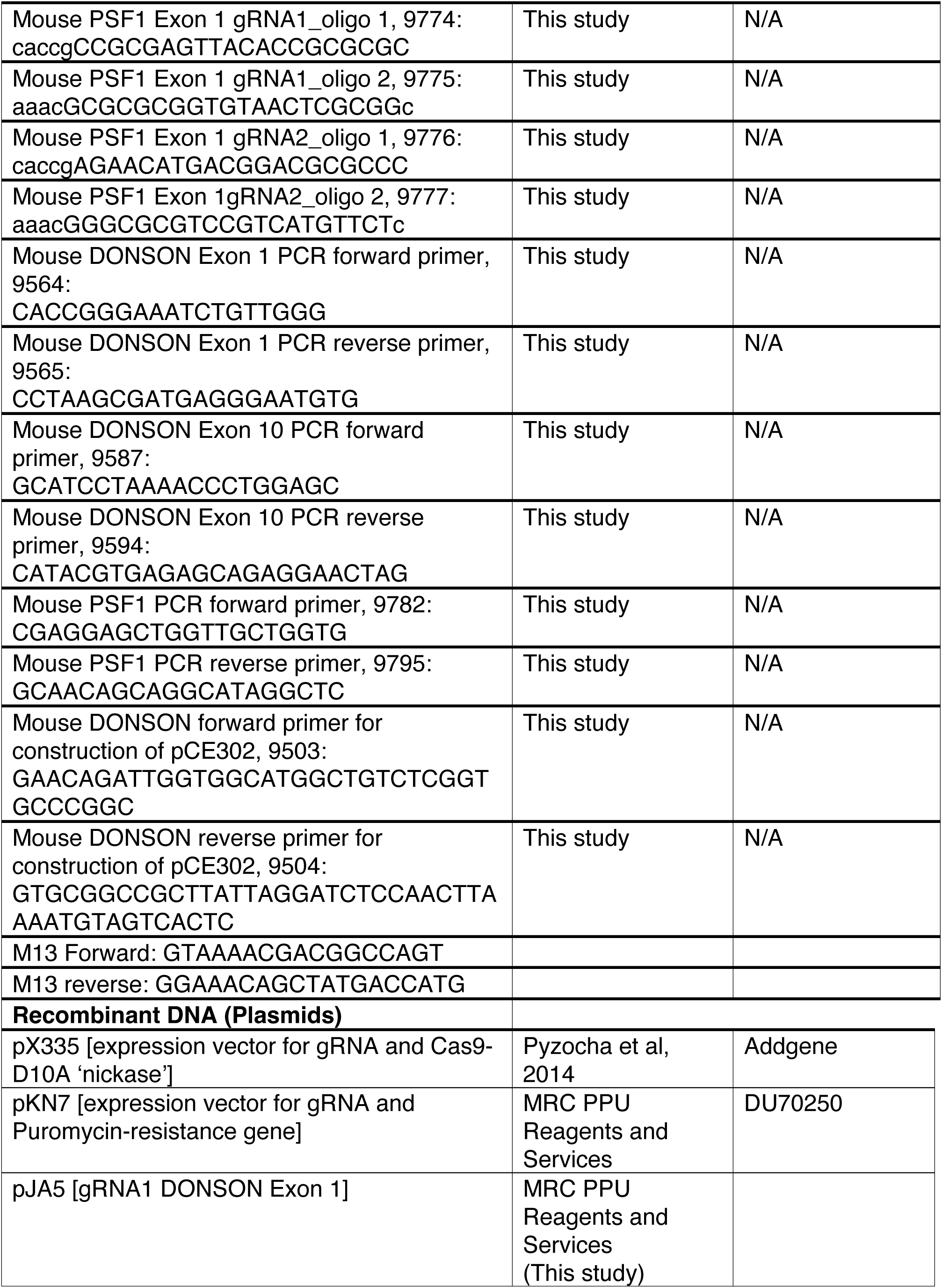
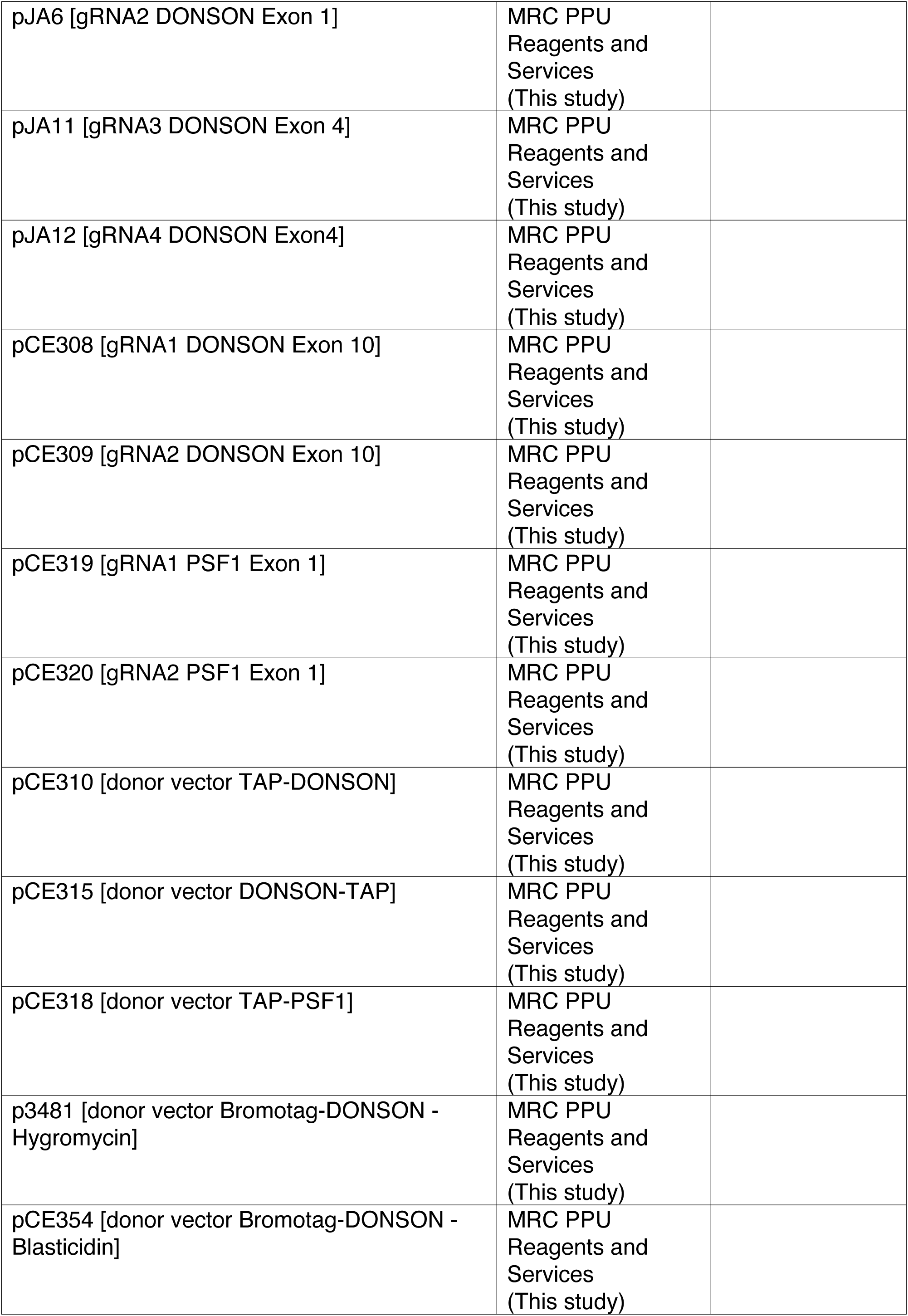
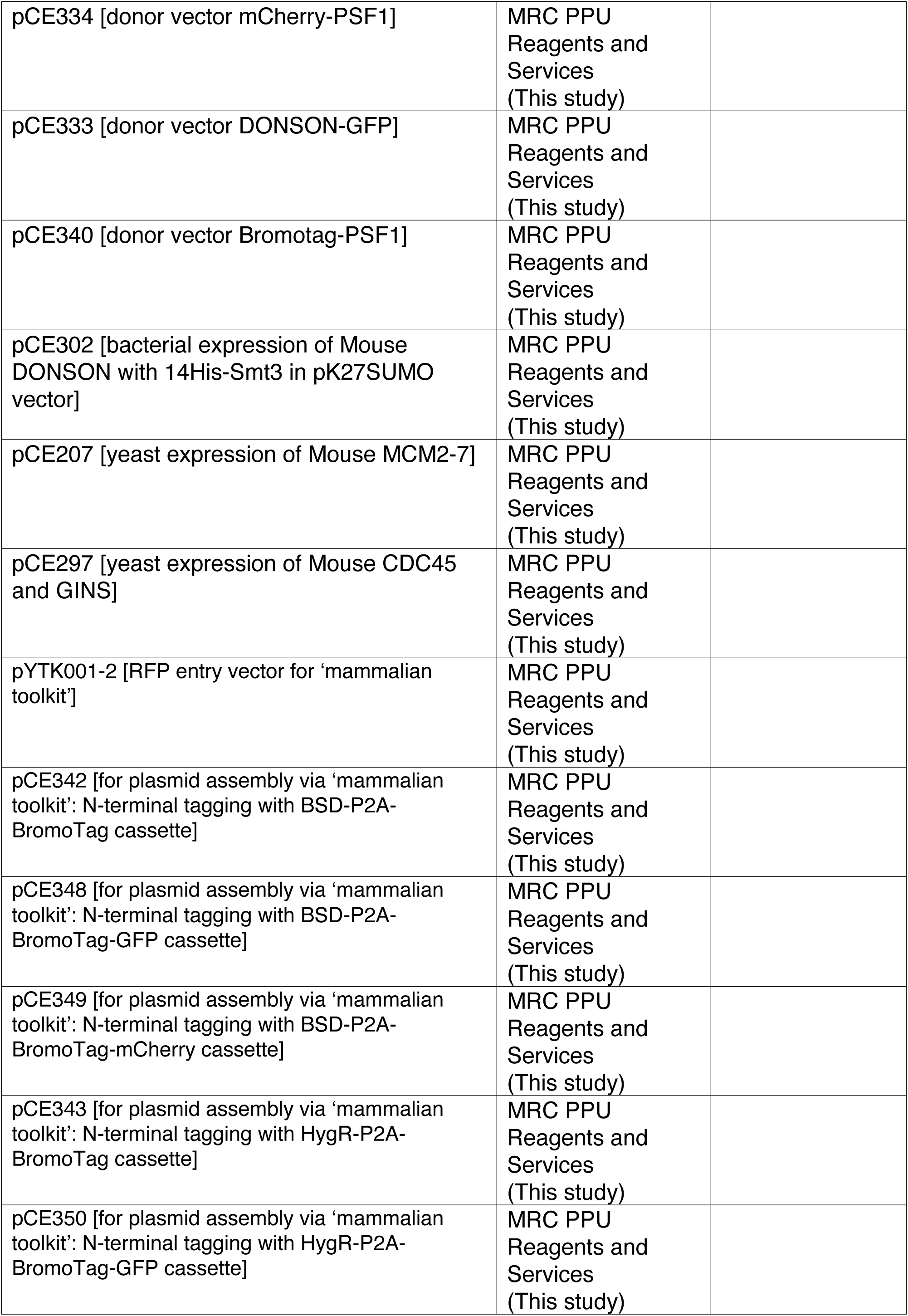
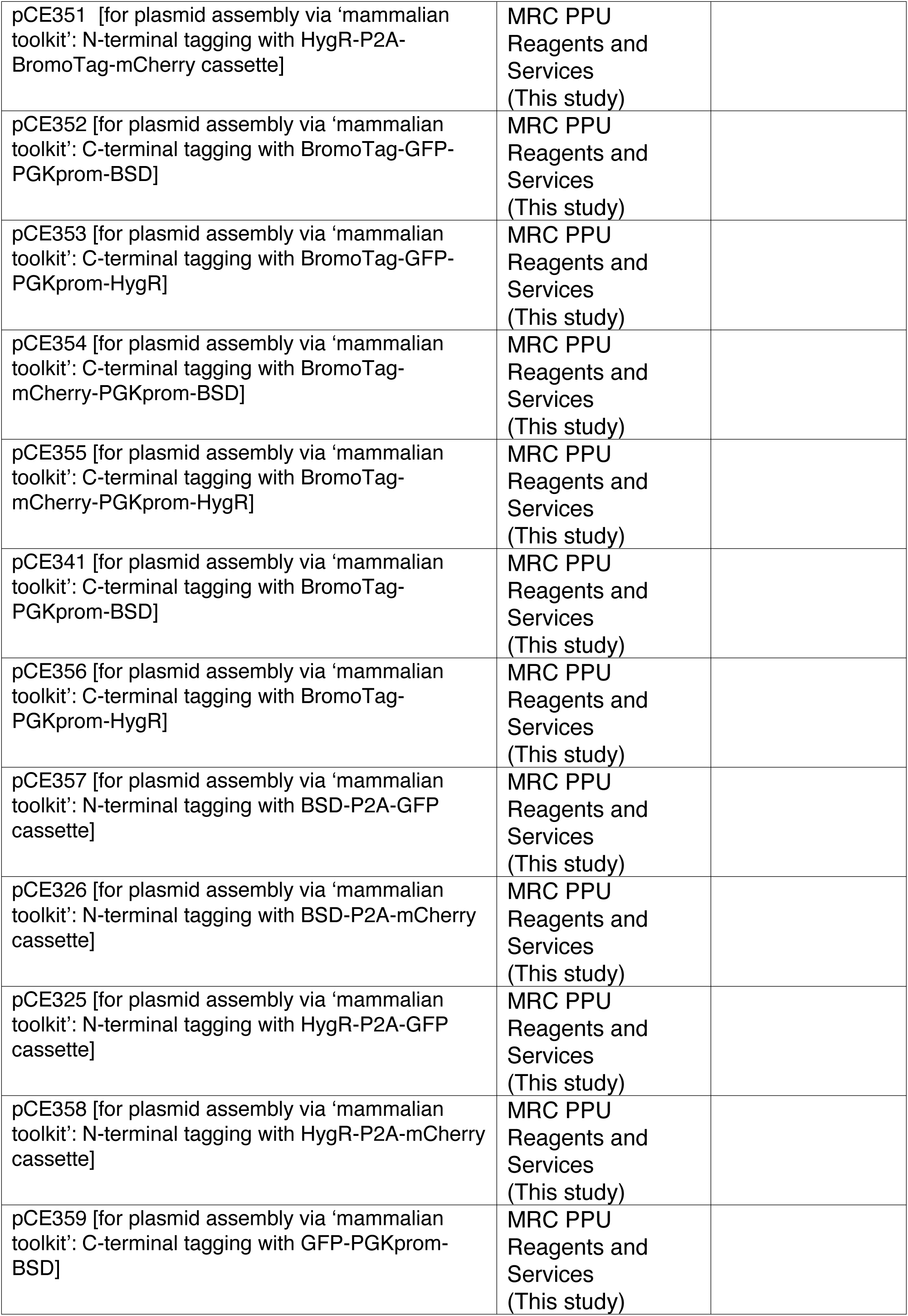
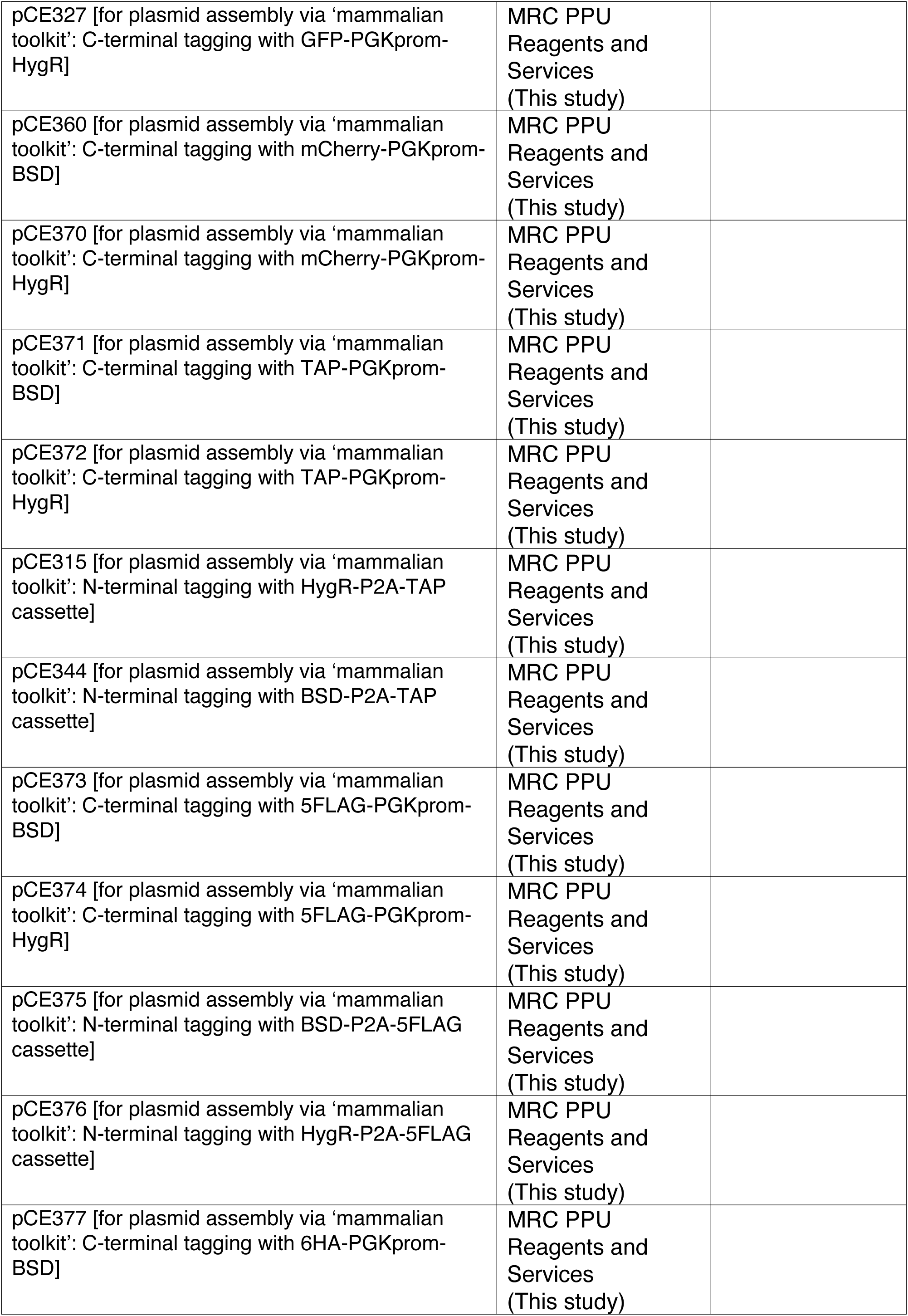
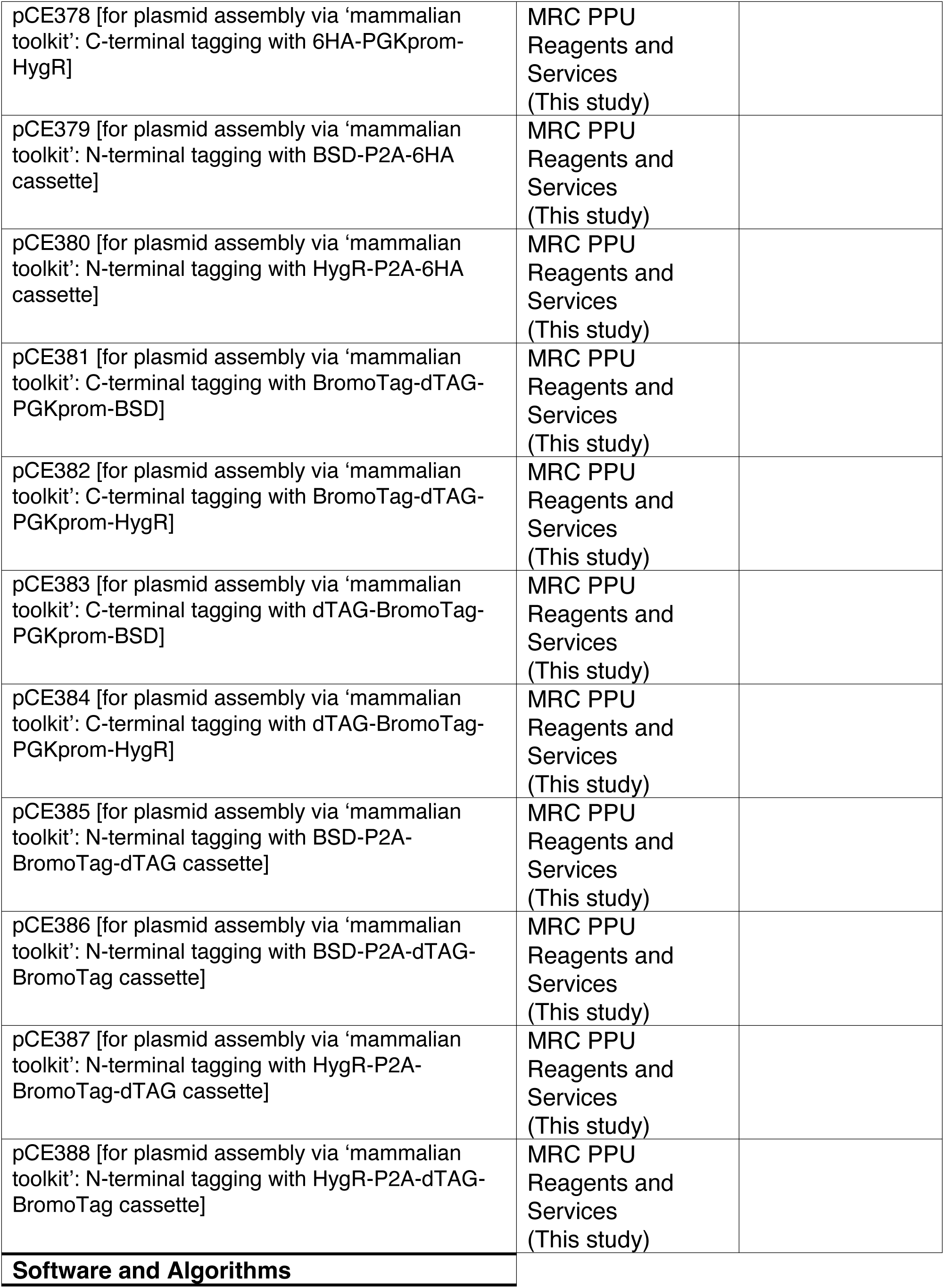
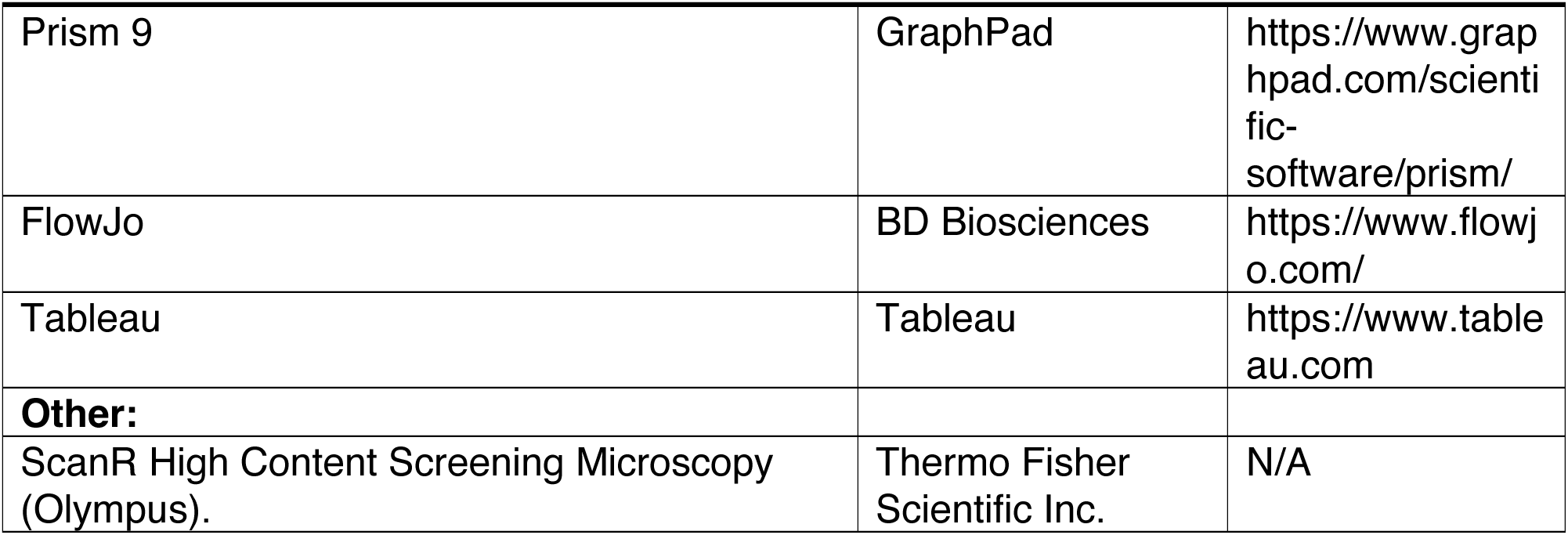
Reagents and resources used in this study.

